# Developmental growth rates adapt to enable self-correction of organ morphology after injury

**DOI:** 10.64898/2026.07.31.741988

**Authors:** S Gunnan, M Kotz, L Ribas, VA Kini, S Jay, S Kaufmann, E Angermann, N Berbee, C Perugini, BM Friedrich, R Mateus

**Affiliations:** Max Planck Institute of Molecular Cell Biology and Genetics; 01307 Dresden, Germany; Cluster of Excellence Physics of Life; Technische Universität Dresden; 01062 Dresden, Germany

**Keywords:** Organ size and shape, growth rate, robustness, feedback control, injury, morphogens, zebrafish pectoral fin

## Abstract

Proportional organ growth requires tissue-level control of cell behavior. Yet the cues that adapt growth rates, especially after organ injury, remain unclear. Here, we uncover that developing organs are robust to extensive damage, triggering injury-specific mechanisms sufficient to override genetically-encoded size defects. Using precision microsurgery in developing zebrafish pectoral fins, we find that injury drives growth asynchronously across tissues and fin axes, by increasing proliferation and extracellular spacing. Growth compensation scales with the amount of tissue lost, restoring size and structure without compromising developmental timing. A feedback-control model captures these dynamics, suggesting that growth rates are regulated toward an organ-specific target area. At the molecular scale, injury signals bypass developmentally-regulated BMP gradient scaling. Injury instead activates *de novo* BMP signaling, which supports growth adaptation and rescues wildtype fin size in developmentally small mutants. Our findings identify an injury-dependent compensatory growth mechanism that resets developmental organ size, ensuring functional organ recovery.

## Introduction

After initial morphogenesis, organ dimensions robustly scale with animal size, establishing a stereotyped body plan^1,2^. This proportional growth requires precise control of tissue growth rates, as imbalances impair organ function. Yet, during organ development, growth is highly dynamic, with growth rates often changing in time and space. To regulate this process, cells interpret positional information cues within the organ^3,4^, sensing both their location and the organ’s overall size, and tuning their proliferative behavior accordingly. However, it remains unclear how these signals dynamically adjust growth rates to instruct patterned and proportionate organ growth, especially when challenged by genetic and environmental stressors, including severe organ injury.

Embryonic development is remarkably resilient to perturbations^5–7^. This developmental robustness is known to emerge from network architectures, *e.g.* non-linear feedback loops, which reduce the propagation of variation and noise from upstream inputs to downstream patterning^8,9^. This can be mediated at the molecular scale by chaperone-mediated proteostasis, shown to buffer against developmental variation and cytotoxic stress^10–15^; at the cellular level, compensatory proliferation and cell type plasticity are known to respond to apoptosis-triggered cell loss^16–18^. While these compensatory mechanisms are understudied, they require a tight link to those providing tissue-level positional information to safeguard functional organ morphogenesis. For example, morphogen production, coupled with extracellular ligand diffusion and degradation, can produce spatial concentration gradients informing cells about their position in developmental fields^4,19,20^.

Across species, Bone Morphogenetic Proteins (BMP) form concentration-dependent signaling gradients^21–25^. Crucially, some of these signaling gradients can scale with tissue size^21,25–29^, a process thought to be regulated by feedback-loops^9,21,30^. Scaling dynamically adjusts the gradient’s signaling range as the tissue grows, thus activating gene expression in a concentration-dependent manner, while tuning organ growth control^21,26,27^. Such organ-wide gradient scaling often requires direct interactions between BMP ligands and heparan-sulfate proteoglycans or glycoproteins^21,31–35^, which in turn govern the morphogen’s extracellular propagation, lifetime, retention, and receptor access across large length scales^31,35^.

Intriguingly, although developmental processes and injury responses entail distinct biological programs, these can become linked in regeneration-competent organs^36–39^. Similar to development, regenerative growth in adult animals is tightly regulated to regain organ size, structure and function. To achieve this, epimorphic regenerating organs, e.g. adult zebrafish fins, rely on positional memory mechanisms, where cells retain positional identity, and respond with position-dependent growth rates upon injury^36,40–44^. Adapting these rates proportionally to the amount of tissue lost ensures that organs resiliently regenerate their pre-injury size and shape. Notably, this recovery time is largely independent of the amputation size.

Despite these advances, developmental and regenerative growth have rarely been examined within a common quantitative framework, and spatially resolved comparisons under matched perturbations remain scarce. It is therefore unclear whether the two share control mechanisms, or whether one can substitute for the other. On one hand, processes inherent to embryonic developmental robustness might deploy compensatory growth mechanisms to buffer for injury-induced stress^16,17,45,46^. On the other hand, injury-induced regenerative programs may overcome developmental defects^47,48^. We therefore asked, how does injury affect growth rates in a developing vertebrate organ? How do those dynamics connect with developmentally-regulated positional information cues, *e.g.* morphogen gradients?

Here, we established precision microsurgery in developing zebrafish pectoral fins, using ultraviolet (UV)-laser microdissection to reproducibly inflict large-scale fin lesions. This allowed us to spatiotemporally map the cellular and tissue dynamics of organ growth adaptation under diverse injury scenarios, comparing those to unperturbed fins. We found that by activating organ-wide cell proliferation and increasing extracellular space, injury leads to compensatory fin growth that increases with the amount of tissue lost, restoring organ size and structure without compromising developmental timing. To rationalize how such growth rates are adapted, we developed a feedback-control model which suggests that fin size is sensed, integrated and adjusted towards an organ-specific target area. At the molecular level, we unexpectedly found that injury triggers specific signals capable of bypassing existing developmental growth regulation, such as the BMP signaling gradients deployed in the pectoral fin. Injury is thus sufficient to rescue fin growth in BMP gradient scaling mutants, restoring size-defective fins to the wildtype target area. Our findings uncover an injury-dependent compensatory growth rate mechanism that resets organ size, buffering against existing developmentally encoded genetic defects.

## Results

### Larval pectoral fins reproducibly resolve large injuries as development unfolds

To systematically and quantitatively investigate the dynamics of adaptive organ growth, we developed a UV-laser microdissection protocol that enables precise removal of a large distal portion of the larval zebrafish pectoral fin, the homolog of the tetrapod forelimbs^49,50^. Without compromising the underlying yolk sac, this method resects the planar, multilayered fin across its proximal-distal axis; when applied to cut 30% of the distal fin at 48 hours post fertilization (hpf), it effectively cuts through five tissue layers^51,52^: the four epithelial layers that comprise the distal fin fold and the endoskeletal disc, i.e. fin’s cartilaginous layer, leaving the proximal fin muscle layers untouched (Fig. 1A, Methods).

**Figure 1.**
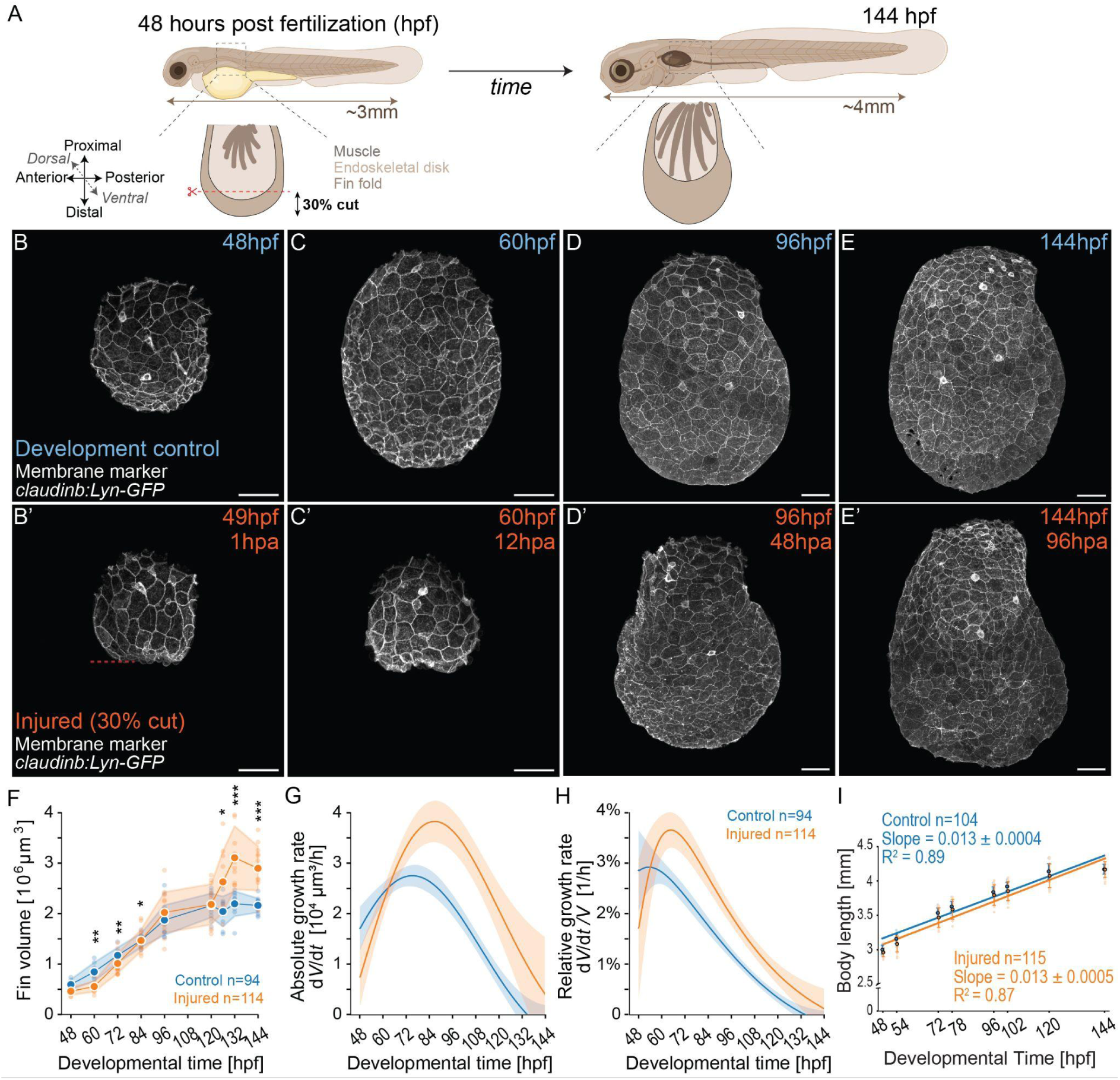
Injury alters pectoral fin developmental growth dynamics, enabling complete organ repair. **(A)** Pectoral fin injury assay applied to zebrafish larvae at 48 hours post fertilization (hpf), highlighting fin developmental axes, region of amputation (red dash) and tissues affected. (**B-E)** Developing control pectoral fins in transgenics labeling epithelial cell membranes (*claudinb:*Lyn-GFP) at 48 (**B**), 60 (**C**), 96 (**D**) and 144 (**E**) hpf. **(B’-E’)** Injured pectoral fins in transgenics labeling epithelial cell membranes (*claudinb*:Lyn-GFP) at 49 (**B’**), 60 (**C’**), 96 (**C’**), and 144 (**D’**) hpf; corresponding hours post amputation (hpa) are marked. Red dash, amputation site. For all images: anterior, left; distal down. Scale bars: 50 μm. **(F)** Average fin volume, *V,* as a function of developmental time in control (blue) *versus* injured (orange) pectoral fins. **(G-H)** Absolute fin growth rate, *dV/dt* (**G**), and relative fin growth rate, *(dV/dt) **/** V* (**H**), as a function of time. Computed from volume *V*(*t*) in F, using smoothing (with temporal resolution 50h, see Methods). **(I)** Larval body length as a function of time, in injured (orange) and control (blue) pectoral fins. Lines represent linear fits with respective goodness of fit, R^2^. Note similarity of slope (i.e. growth rate) between conditions. In F-H: mean values connected by solid lines; shading indicates ±SD. *n*, number of fins (F-H) or larvae (I); 7-17 per timepoint. All statistics: \**p*≤0.05, \*\**p*≤0.01, \*\*\**p*≤0.001; two-tailed, unpaired, non-parametric Mann-Whitney tests.

We first quantified the pectoral fin’s postinjury volume dynamics, by employing our microdissection assay coupled to live imaging of an epithelial membrane reporter line (Tg(−8.0cldnb:lynGFP)^zf106^, i.e., claudinb:Lyn-GFP)^53^. By measuring the fin’s volume at defined time points, in injured and age-matched developmental controls, we identified that this organ reproducibly recovers its functional morphology within four days (Fig. 1A-E). Interestingly, we found that the fin’s volumetric growth dynamics are altered upon injury (Fig. 1F): in the first 12 hours post amputation (hpa), between 48 and 60 hpf, fin volume increases only marginally. From this time onwards, the fin’s mean volume consistently rises and within two days after injury (96 hpf, 48 hpa), the volume of injured fins has caught up with that of uninjured fins. Intriguingly, the volume of fins subjected to injury surpasses that of corresponding uncut fins by 132 hpf (84 hpa, 41.6%), a trait that persists in recovered fins, four days after injury (144 hpf, 96 hpa). In contrast, the average volume of fins undergoing solely development increases monotonically until 96 hpf; after this time, growth arrests, and consequently their volume plateaus at ≈1.8 x10^6^ µm^3^. This growth pattern is temporally consistent with former observations where BMP signaling gradients present from 48 to 78 hpf in the pectoral fin play a major role in regulating its developmental size^21^.

Next, by estimating the absolute and relative growth rates in injured and control fins, we quantified how much these two growth contexts differ from one another, pinpointing when the injured fins adapt their growth dynamics, as development proceeds. While in the initial 12 hours postinjury, the damaged fins’ growth rate is lower than those undamaged, consistent with a wound healing phase^36,54^, after this, the injured fins’ growth rate is systematically higher than in unperturbed controls (Fig. 1G-H). In contrast, these unperturbed fins display growth rates that decrease to zero, in line with the observed volume growth plateau (Fig. 1F) and previous reports of pectoral fin development^21^.

We conclude that larval pectoral fins successfully recover from extensive injuries inflicted at two days post fertilization. Remarkably, as this organ’s repair process occurs, fin development is not interrupted, and the zebrafish larva continues to increase its body length on average by ∼33% between 48 and 144 hpf (Fig.1I). Hence, in contrast to other developing systems^55^, recovery from injury in the zebrafish larval pectoral fin does not exclusively restore its damaged tissues, after which organismal development is resumed. Instead, the injured pectoral fin dynamically adapts its growth rate, whereby this organ ‘catches up’ with developmental size and architecture as it heals, resulting in a functional fin by 144 hpf.

### Global proliferation and extracellular volume restore tissue proportions in injured fins

Our comparative measurements of the larval pectoral fin volume dynamics led to two central questions underlying the robustness of our system to perturbation: Which fin tissues respond to injury? What cellular behaviors underlie the observed fin growth rates? To address the first question, we imaged transgenic reporter lines labeling specific tissues of the larval pectoral fin, as well as this organ’s extracellular space (Fig. 2A-E, Methods), in uncut and injured conditions.

**Figure 2.**
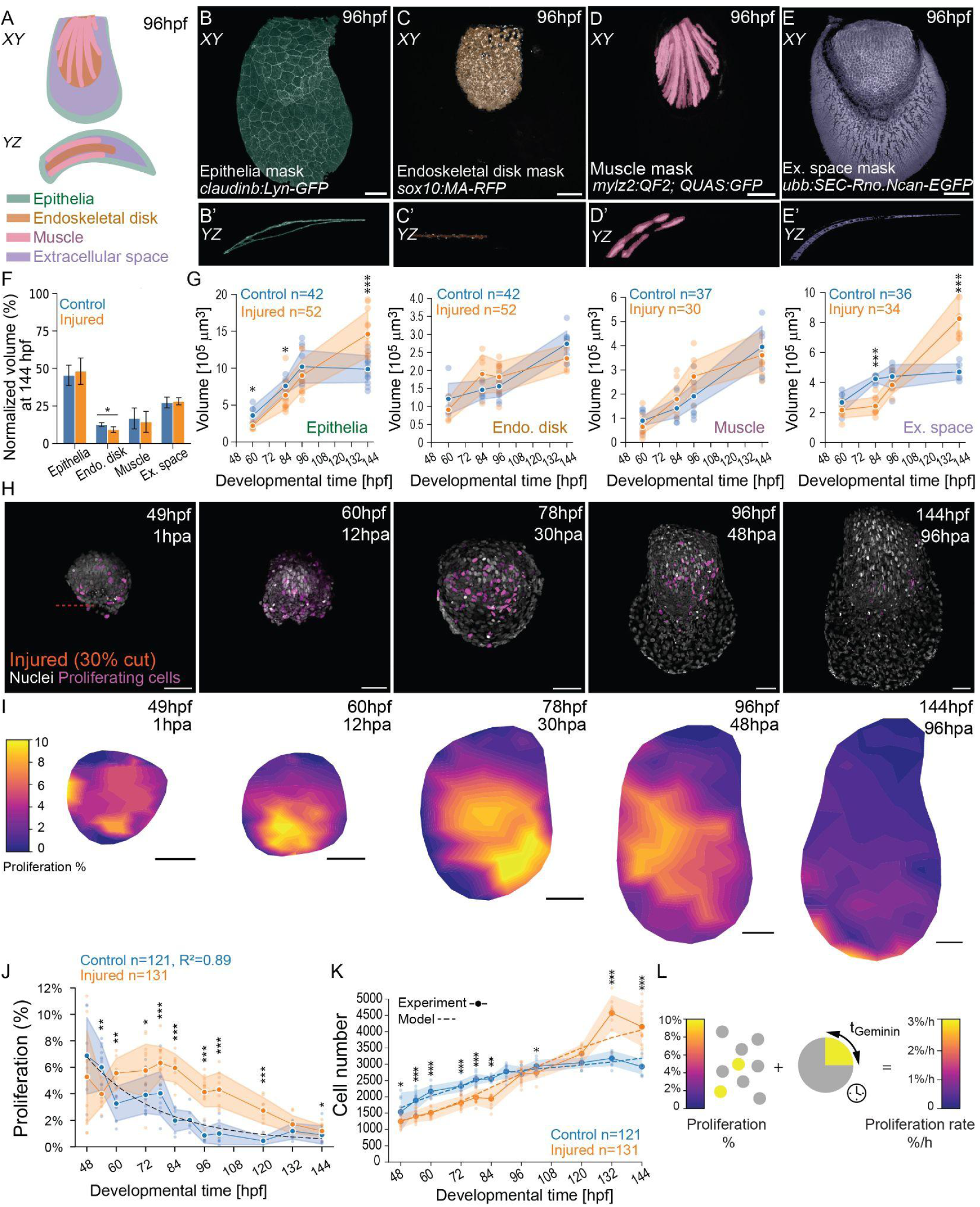
Tissue-specific growth and cell proliferation dynamics compensate for injury. **(A)** Schematics highlighting pectoral fin tissue organization at 96 hpf, in *XY* and *YZ* views. **(B-E)** Pectoral fins from transgenic reporter lines labeling the epithelia (**B**), endoskeletal disk (**C**), muscle (**D**), and extracellular space (**E**), with respective *YZ* views (**B’-E**’), at 96 hpf. Note that segmentation masks are overlaid in each image following the legend in A. **(F)** Comparison of relative volume (%) of each tissue of the fin normalized by respective fin volumes at 144 hpf, in control (blue) versus injured (orange) fins. **(G)** Volume comparisons in control (blue) versus injured (orange) pectoral fins’ epithelia, endoskeletal disk, muscle, and extracellular space, between 60-96 hpf. **(H)** Pectoral fins from transgenics labeling all nuclei (H2B-mCherry, white) and proliferating cells (mAG-zGeminin^+^, magenta), upon injury. Red dash, amputation site. **(I)** Spatial density maps of proliferation in injured fins, from 49 hpf (1 hpa) to 144 hpf (96 hpa). Color code shows the percentage of zGeminin^+^ nuclei. zGeminin^+^ nuclei were projected onto fin midsurface and averaged over *n* = 8-17 fins per condition (Methods). **(J)** Percentage of proliferative (zGeminin^+^) per total cells, in control (blue) versus injured (orange) fins, from 48 to 144 hpf. Black dash, exponential fit to individual points of the control dataset, with goodness of fit, R^2^. **(K)** Average cell number in control versus injured fins can be explained by a growth model (dashed lines) that integrates the proliferation in time, assuming a constant proportionality factor of 3.25 h between proliferation rate and the fraction of proliferating cells, indicative of the duration of the zGeminin^+^ phase of the cell cycle (Fig. S1E-G). **(L)** The proliferation amount (%) can be converted into a proliferation rate (%/h). For all images: anterior, left; distal down. Scale bars: 50 μm. *n*, number of fins. In G, J-K: mean values connected by solid lines; shading indicates ±SD. All statistics: \**p*≤0.05, \*\**p*≤0.01, \*\*\**p*≤0.001; two-tailed, unpaired, non-parametric Mann-Whitney tests.

By segmenting each fin tissue’s volume and quantifying those dynamics between 60 and 144 hpf, we first confirmed that when unperturbed, each fin tissue develops asynchronously (Fig. 2G, blue)^52,56,57^. In doing so, our analysis exposed the relative contributions of each tissue to the overall fin size (Fig. S1A-B). For example, at 60 hpf, the developing pectoral fin is composed of approximately 43% epithelia, 14% cartilage (endoskeletal disk), 8% muscle, and 33% extracellular space (Fig. S1A-B). Upon injury, we unexpectedly found that each of these tissues regrows with different dynamics compared to uninjured controls, highlighting the presence of tissue-specific adaptive growth (Fig. 2G, orange). In the first two days postinjury (up to 96 hpf, 48hpa), injured tissues like the cartilage or the muscle grow more than uninjured counterparts (Fig. 2G, Fig. S1A-B). After this phase, epithelial regrowth as well as an increase in extracellular spacing surpass those of uninjured fins (Fig. 2G, Fig. S1A-B), justifying the previously measured volume increase in injured fins (Fig. 1F). Despite the asynchronous tissue growth and irrespective of having damage inflicted, by 144 hpf, we observe that fin tissues show similar volumes relative to the respective total volume (Fig. 2F, Fig. S1A-B). We conclude that the overall organ growth dynamics observed result from a concerted increase in volume from each of the individual fin tissues as well as a non-negligible contribution from extracellular volume.

The observed restoration of tissue-specific proportions upon pectoral fin injury showcases a striking example of embryonic development buffering against extensive cell loss. We therefore asked whether this resilience relies on classical modes of regenerative growth. Interestingly, we found no evidence of epimorphosis^36,38^: injured fins did not display single-cell migration from any of the analysed tissues into the wound site, loss of lineage-specific reporter signal at particular timepoints (indicative of de-differentiation or transdifferentiation processes), or contribution of the analysed tissues into a blastema-like structure (Fig. S1A-B). To then understand whether the observed fin growth rates could be accounted for by a dynamic compensation of cell division, we set out to quantify the fraction of proliferative cells in developing *versus* injured fins, at defined time points. For this, we applied our injury assay coupled with live imaging of transgenic reporter lines labeling all nuclei (Tg(Xla.Eef1a1:hist2h2l-mCherry)^zf502Tg^, *i.e.* H2B-mCherry)^58^ as well as proliferative nuclei (S/G2/M FUCCI marker, Tg(EF1a:mAG-zGem(1/100))^rw041^, *i.e.* mAG-zGeminin^+^)^59^, followed by 3D nuclei segmentation and quantification (Fig. 2H, Fig. S1C, Methods).

By calculating the percentage of proliferating cells over time, we found that the observed fin growth rates appear largely dependent on the dynamics of cell division, regardless of the presence of injury. First, we confirmed that developing fins have an exponentially decaying proliferation rate (Fig. 2J, blue; Fig. S1C)^21^. In contrast, age-matched injured fins display proliferation dynamics characteristic of regeneration-competent systems (Fig. 2J, orange)^36^: in the initial hours following damage, 1-6 hpa (49-54 hpf), the proliferation rate is lower than in uninjured fins; then, by 12 hpa (60 hpf), the fraction of proliferating cells in injured fins increases significantly, remaining consistently higher than that of developing fins until 84 hpa (132 hpf). Furthermore, quantitative spatial analysis of cell proliferation showed that proliferation levels are mostly homogeneous throughout the fin (Fig. 2I), a feature observed also in uninjured fins (Fig. S1D)^21^.

Finally, rationalizing that the observed proliferation dynamics should be reflected in increased cell numbers, we then compared the total number *N*(*t*) of fin cells as a function of time, with the prediction of a simple growth model that integrates the time-dependent mean proliferation rate for both uncut and injury conditions (Fig. 2K, Methods). Specifically, we use the total number of segmented nuclei as proxy for the total cell number *N*(*t*), and the fraction of proliferating cells as proxy for the relative proliferation rate, (d*N*/d*t*) / *N*. To convert the fraction of proliferating cells (with units %) to a rate (with units %/h), we divide this fraction by a putative duration of the proliferative phase of the cell cycle, *i.e.*, the zGeminin^+^ phase (Methods). We determined this value to be ∼3.25 h in both injured and control fins by a fit that compares the measured cell number *N*(*t*) and the expected cell number reconstructed from the proliferation rate. The estimated zGeminin^+^ duration is comparable, albeit shorter than direct measurements of the S/G2/M phase in developing pectoral fins (5.4 ± 1.2h, Fig. S1E-G).

The close match between the measured and predicted number of cells, using the simple assumption of a constant and spatially homogeneous duration of the zGeminin^+^ phase in both injured and control fins, validates our proliferation analysis (Fig. 2J-K). Furthermore, it calibrates a suggested conversion of the measured fraction of zGeminin^+^ cells into a true proliferation rate, now possible to be interpreted in developmental time and fin space (Figs. 2I-L; Fig. S1D). Together, our results demonstrate that while injured pectoral fin tissues regrow asynchronously (Fig. 2F-G), their recovery is jointly driven by an adaptive response in cell proliferation (Fig. 2J-K) and an increase in extracellular spacing (Fig. 2F-G), accounting for the measured fin volumes.

### Pectoral fins recover their morphological proportions within two days of injury

Our results suggest that the observed injury-induced changes in developmental growth rates are a consequence of elevated cell proliferation and expanded extracellular space across the pectoral fin. Together, these collective behaviors promote the fin’s size self-correction upon injury. But how do these growth rates affect shape recovery?

To quantify fin shape over time, we computed a two-dimensional midsurface (Fig. 3A, Fig. S2, Methods) that allowed unprecedented rigorous measurement of the lengths of each of the fin’s curved developmental axes in three space dimensions, across our previously acquired volume datasets (Fig. 1F). In injured fins, we observe that the lengths along the proximal-distal (PD) and anterior-posterior (AP) axes have approximately reached those of uninjured controls by 96 hpf (48 hpa)(Fig. 3B). Unexpectedly, the length along the dorso-ventral (DV) axis significantly increases post-injury, with its average values surpassing those of uninjured fins by 60 hpf (12 hpa), and remaining higher than uncut controls by 144 hpf (96 hpa, 44.5 µm cut *vs.* 40.8 µm uncut, Fig. 3B). To confirm that the length increase along the DV axis upon injury was not a local effect, we also computed the overall fin thickness (Fig. S3A, Methods), which displayed a similar growth trend. Therefore, injury uncouples growth across the pectoral fin developmental axes, leading to an increase in fin thickness, which accounts for the growth ‘overshoot’ in volume observed in the later stages of fin recovery (Fig. 1F, orange).

**Figure 3.**
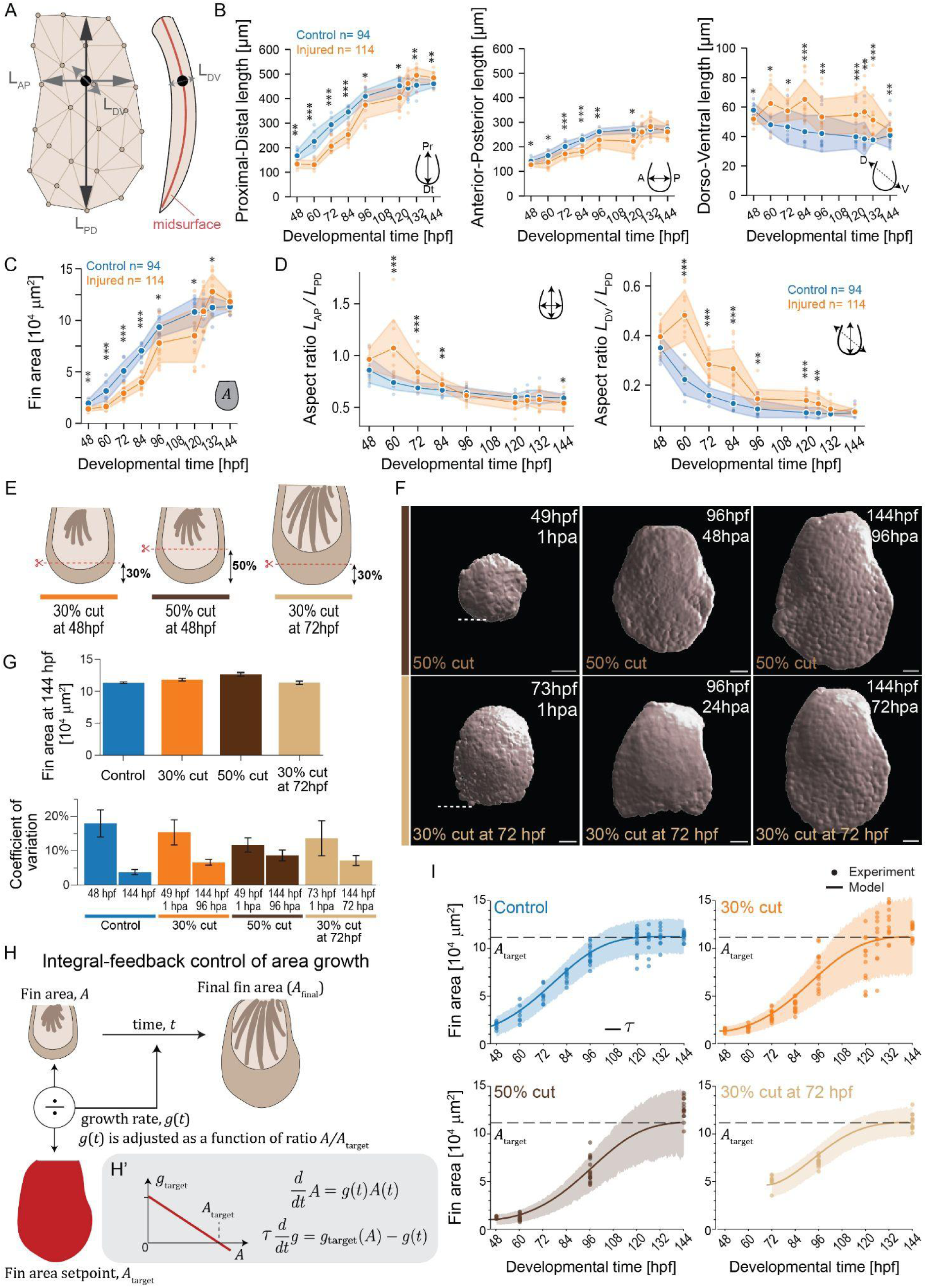
Adaptation in growth rate is compatible with a controlled setpoint in fin area. **(A)** Two-dimensional midsurface (red) used to compute fin surface area and lengths along the three fin axes (*L*_PD,_ *L*_AP,_ *L*_DV_) from volumetric 3D masks. The coordinate axes used for length measurements all pass through a reference point located at a relative position of 40% of the proximal-distal axis and 50% of the anterior-posterior axis; the dorso-ventral axis length is computed at this reference point along the normal direction. **(B)** *From left to right*, average length along proximal-distal, anterior-posterior or dorso-ventral fin axes (defined in A) as a function of time, in control (blue) or injured (orange) fins. **(C)** Fin area, computed as the area of the fin midsurface, as a function of time in control (blue) or injured (orange) fins. **(D)** Fin aspect ratio as a function of time, between the anterior-posterior and proximal-distal axes (*left*), and dorso-ventral and proximal-distal axes (*right*), in control (blue) or injured (orange) fins. Aspect ratios close to one indicate a near circular cross-section as, e.g., opposed to an elongated one. **(E)** Additional injury scenarios tested: distal cut of 50% at 48 hpf (*middle*, dark brown), and distal cut of 30% at 72 hpf (*right*, beige). **(F)** 3D surface masks of pectoral fins live imaged for the two additional injury scenarios from E: 50% cut at 48 hpf, and 30% cut at 72 hpf. Note that masks were obtained from nuclei signals (H2B-mCherry, Methods). White dash, amputation site. **(G)** Comparisons of fin area at 144 hpf (*top*), and sample standard deviation normalized by sample mean (i.e., coefficient-of-variation, *bottom*) at 48 and 144 hpf, for control (blue) and the three tested injury scenarios. Irrespective of perturbation, the fin area converges to an approximately constant value with simultaneous reduction in relative variation. **(H)** Growth model with a dynamic growth rate *g(t)* regulated according to an area setpoint. **(H’)** Model equations. Note that *g*(*t*) is regulated towards a target value *g*_target_(*A*) that decreases with fin area *A*, becoming zero when *A* equals target size, *A*_target_. **(I)** Comparison between fin area obtained by experimental data (dots, from C and F) and model (line). Note that the model can be simultaneously fit to the four growth scenarios, only assuming different initial conditions (Methods, Supplementary Theory Notes). For all images: anterior, left; distal down. Scale bars: 50 μm. In B-D: mean values connected by solid lines; shading indicates ±SD. *n*, number of fins, 7-17 per timepoint. All statistics: \**p*≤0.05, \*\**p*≤0.01, \*\*\**p*≤0.001; two-tailed, unpaired, non-parametric Mann-Whitney tests.

Notably, the proximal-distal and anterior-posterior axes show similar length dynamics regardless of injury, despite differing in magnitude (Fig. 3B *left* vs. *middle*), a feature also captured by the fin area measurements (Fig. 3C). To further understand how growth along the different organ axes contributes to fin shape, we computed aspect ratios between the lengths of the anterior-posterior and proximal-distal axes (*L*_AP_ / *L*_PD_), as well as between dorsal-ventral and proximal-distal axes (*L*_DV_ / *L*_PD_), over time and across our datasets (Fig. 3D). The analysis revealed that the injured fins are rounder than uninjured controls in the first 12 hours post amputation (60 hpf), after which the fin’s characteristic elongated shape^21,52^, i.e. a longer proximal-distal axis, starts to be regained (Fig. 3D, *left*). This is reflected in the fin’s anisotropic growth rate, where the proximal-distal axis presents larger growth rates than the anterior-posterior axis, regardless of injury (Fig. S3B). Interestingly, by 48 hpa (96 hpf), the fin’s area aspect ratio (i.e. (*L*_AP_ / *L*_PD_)) is fully restored, with the fin displaying 2D proportions similar to those of developmental controls (Fig. 3D, *left*). Similarly, when considering the relation between dorsal-ventral and proximal-distal axes (*L*_DV_ / *L*_PD_), we detected that injured fins have their proportions altered until 48 hpa (96 hpf), instead of progressively becoming thinner and elongated as in uncut controls (Fig. 3D, *right*). We conclude that while growth along the dorso-ventral axis appears uncoupled from the other two fin axes (Fig. 3B), fin proportions are remarkably recovered within two days after injury (Fig. 3D). This points towards an active mechanism regulating fin size and shape to fast-track organ functionality.

### Fin growth dynamics converge on a single target area, regardless of injury

Our findings exemplify how developing organs are resilient to large internal and external perturbations^6^. By actively modulating their growth rates, cells compensate for tissue loss while restoring age-matched size and pattern, buffering stress, while informed by active positional information mechanisms^6,20^. In contrast, adaptive, position-dependent growth rates are a hallmark of adult regeneration-competent organs^36,40,41,43,44,60^. Here, organs long devoid of positional information-conferring mechanisms, retain a positional memory that, when triggered by injury, leads to specific cellular source activation, restoring only the parts lost^42,60,61^. Due to the similarities in growth adaptation dynamics, we asked whether injury in developing organs can trigger features traditionally associated with positional memory. To fill this gap, we extended our experimental assay to explore how differences in proximo-distal injury position could be linked to an adaptation of the fin growth rates, and whether this is specific to a developmental time-window (Fig. 3E).

Remarkably, regardless of increasing the injury size at 48 hpf or applying it one day later (Fig. 3E-F), the fins recovered their volume, area and thickness to values within error of the developmental uncut controls, by 96 hpa (144 hpf) (Fig. 3G,I; Fig. S3D). Because recovery is completed within the same time window, larger amputations must be compensated by higher growth rates. Thus, the injured larval pectoral fin, like the adult zebrafish or urodele appendages^36,40^, displays growth rates based on the amount of tissue lost, a feature of positional memory inflicted by injury.

Motivated by the apparent coupled growth dynamics along the proximal-distal and anterior-posterior axes occurring irrespective of injury (Figs. 3B-C), we then asked whether the fin area could be used as a size proxy. By 144 hpf, fin area not only reached approximately the same final value across all growth scenarios (Fig. 3G, top), but also showed a 2–3 fold decrease in its coefficient of variation, a measure of relative variability, over time (Fig. 3G, bottom). This is incompatible with a simple timer mechanism that instructs growth for a fixed period of time independent of fin size (Supplemental Theory Notes, Fig. S4, see also ^62,63^). Instead, the observed reduction of the coefficient of variation suggests feedback control. To consider this, we formulated a minimal model of organ growth control, which dynamically adjusts the organ’s current area growth rate in response to the mismatch between the present size and an intrinsically specified target area. In choosing the control variable in our model, we also considered fin volume; however, given that fin growth along the dorso-ventral axis appears decoupled from the in-plane axes (Figs. 3B-C, Fig. S3A), final fin volumes fluctuate more across the different tested conditions, hence the reduction in the coefficient-of-variation was less pronounced (Fig. S3F).

Mathematically, the model implements a variant of integral-feedback control for fin area (Fig. 3H-H’, Supplementary Theory Notes). In short, the fin area *A*, grows with a time-dependent relative growth rate, *g*(*t*). This growth rate is regulated on a time-scale *τ* towards a target growth rate, *g*_target_(*A*), which depends on the ratio between the fin area at a given time *A*(*t*) and the target area, *A*_target_, here chosen as the final area of the fin at 144 hpf. This feedback ensures that *A*(*t*) will always converge to this setpoint *A*_target_, while concomitantly, the growth rate will converge to zero.

Notably, this minimal model is able to fit the fin growth dynamics observed across all experimental scenarios tested, irrespective of injury and assuming only a common, organ-intrinsic set of model parameters (Fig. 3I, Fig. S3E). The only difference between fits is the initial fin area (given that experimental cut conditions differ largely; Figs. 1B, Fig. 3F); the initial growth rate is set to zero in regeneration scenarios, reflecting the growth arrest that occurs during wound healing. From the fit, we obtain a growth control time-scale *τ* = 7.3±1.0 hours, similar to the time after injury without proliferation (Fig. 2J). Importantly, our model suggests that sensing current fin size and consequently adjusting the fin’s growth rate is not instantaneous, but takes time on the order of several hours (Supplementary Theory Notes).

### Injury restores size-defective pectoral fins to wildtype target area

Our mathematical model suggests that injury-specific adaptation of growth rates relies on an intrinsically specified setpoint for fin area. Yet, which molecular mechanisms would implement such a setpoint and is there redundancy of control mechanisms? A common feature of dynamic growth control mechanisms is that these can be enforced by morphogen concentration gradients. Hence, we focused on BMP signaling gradients given their central role in regulating pectoral fin developmental size^21,64^.

Given the proposed BMP-Smoc1 gradient scaling system in the developing pectoral fin^9,21,30^, and recent work showing that direct binding between secreted BMP dimers and Smoc proteins is required for BMP signaling across species (Fig. S5D-E)^65^, we made a specific prediction. Removing the region where Smoc1 is mainly expressed by a 30% cut fin injury (Fig. 4A, Fig. S5A-B) should initially reduce BMP signaling: with less Smoc1 present in the fin, extracellular BMP degradation would increase, or its diffusion decrease. Yet at later times after injury, when growth resumes, we would expect that BMP signaling gradients scale again, with concomitant re-expression of Smoc1.

**Figure 4.**
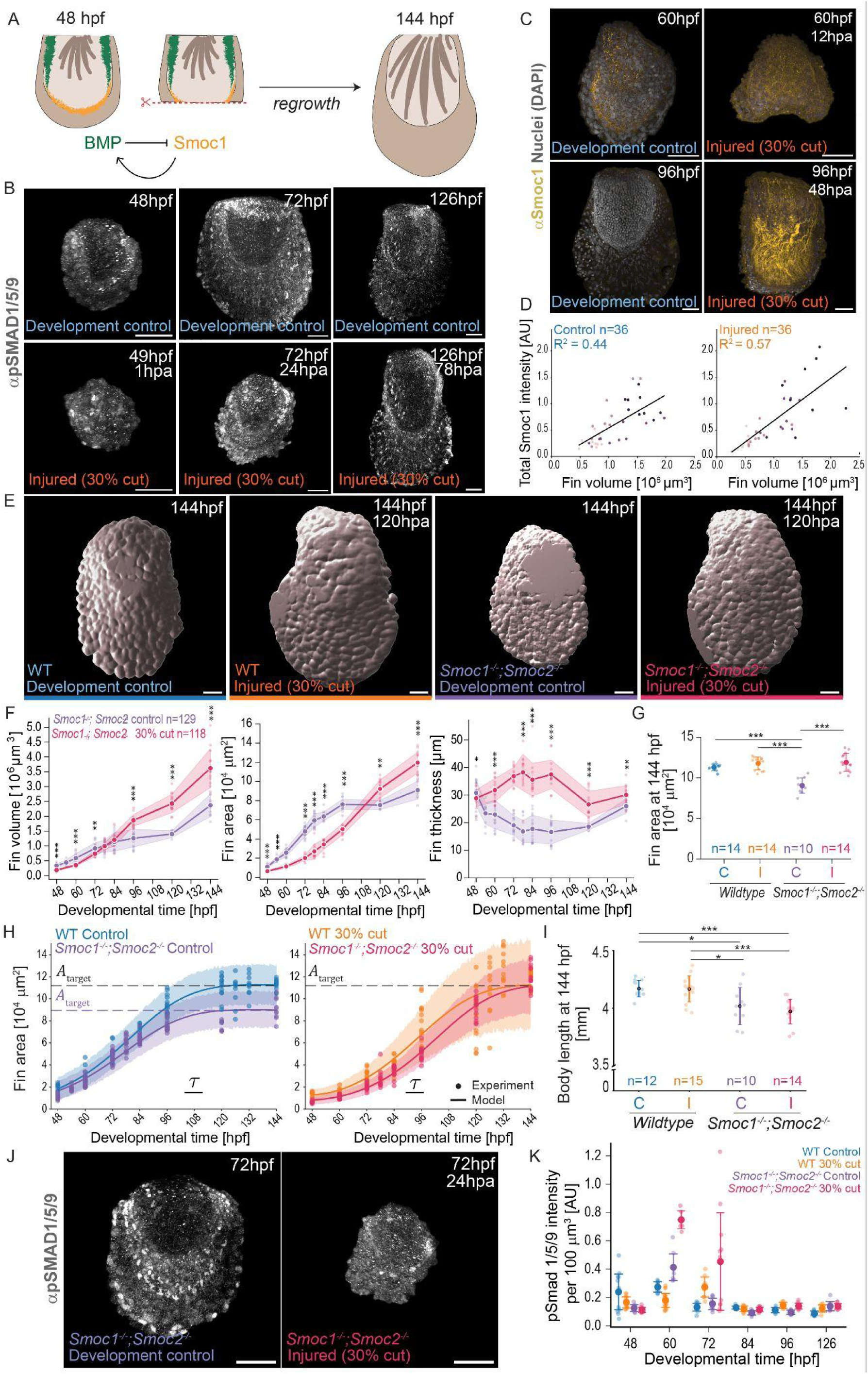
Injury rescues fin growth in BMP signaling gradient-scaling mutants. **(A)** Injury at 48 hpf removes a large part of developmentally-regulated Smoc1 expression in pectoral fins. **(B)** Maximum intensity projections of phosphorylated Smad1/5/9 immunostainings in control and injured fins, at 48 (1 hpa), 72 hpf (24 hpa) and 126 hpf (78 hpa). **(C)** Maximum intensity projections showing Smoc1 expression (yellow) pattern in control and injured fins, at 60 hpf (12 hpa) and 96 hpf (48 hpa). **(D)** Total Smoc1 intensity as a function of fin volume, in control (left) and injured (right) fins, from 60 to 96 hpf (12 to 48 hpa). Lines, linear fits with goodness of fit (R^2^). The darker the dot color, the older the fish. **(E)** *From left to right*, 3D surface masks of pectoral fins live imaged at 144 hpf in different backgrounds: wildtype uncut control, *Smoc1^-/-^;Smoc2^-/-^*uncut control, wildtype 30% cut, and *Smoc1^-/-^;Smoc2^-/-^*30% cut. Note that masks were obtained from nuclei signals (H2B-mCherry, Methods). **(F)** *From left to right*, comparison of fin volume, area or thickness as a function of time between *Smoc1^-/-^;Smoc2^-/-^*uncut control (purple) or injured *Smoc1^-/-^;Smoc2^-/-^*(pink) fins. n = 6-23 fins per condition. **(G)** Comparison of fin area at 144 hpf, between larvae injured with 30% cuts (I) or control (C) pectoral fins, in wildtype or *Smoc1^-/-^;Smoc2^-/-^* backgrounds. **(H)** Comparison of fin area growth between experimental data (dots, from Figs. 3C and 4F) and model (line) for wildtype uncut control (blue), *Smoc1^-/-^;Smoc2^-/-^*uncut control (purple), 30% cut wildtype (orange), 30% cut *Smoc1^-/-^;Smoc2^-/-^*(pink). Model parameters from fit to wildtype data were re-used (Fig. 3I, same parameters only assuming different initial conditions), except for a different target area *A*_target_ for *Smoc1^-/-^;Smoc2^-/-^*uncut control (Methods, Supplementary Theory Notes). **(I)** Comparison of larval body length at 144 hpf, between larvae with injured (I) or control (C) pectoral fins, in wildtype or *Smoc1^-/-^;Smoc2^-/-^* backgrounds. **(J)** Maximum intensity projections of phosphorylated Smad 1/5/9 immunostainings in *Smoc1^-/-^;Smoc2^-/-^* control and injured fins, at 72 hpf (24 hpa). **(K)** Mean intensity of phosphorylated Smad 1/5/9 immunostainings (per 100 µm^3^) between 48 to 126 hpf (1-78 hpa), for control and injured fins, in wildtype and *Smoc1^-/-^;Smoc2^-/-^* backgrounds. For all images: anterior, left; distal down. Scale bars: 50 μm. Shading or bars in all plots, SD. *n*, number of fins (D, G-H, K) or larvae (I); 9-15 per condition/timepoint. Statistics F-G, I: \**p*≤0.05, \*\**p*≤0.01, \*\*\**p*≤0.001; two-tailed, unpaired, non-parametric Mann-Whitney tests.

To test our hypothesis, we applied our injury assay to pectoral fins and quantified BMP signaling. For this, we performed immunostainings against the phosphorylated form of Smad1/5/9 (pSmad1/5/9, a well known BMP signaling readout)^66^ at different timepoints after injury. As expected, right after injury, BMP signaling decreases compared to control fins (Fig. 4B, K). However, at later times after injury, pSmad1/5/9 increases, becoming active not only in the characteristic graded domains in the anterior and posterior sides of the developing fin, but also in an additional region adjacent to the wound (Fig. 4B, K). By quantifying pSmad1/5/9 levels in cut *versus* uncut fins, we observed that at one day after injury (72 hpf, 24 hpa), pSmad1/5/9 levels in injured fins surpasses that of the developing fins (Fig. 4B, K). This boost in BMP signaling is dynamic and reverts back to the baseline of uncut fins by 84 hpf (36 hpa) onwards.

To understand if the increase in BMP signaling upon injury is correlated to Smoc1 upregulation, we then performed comparative immunostainings against Smoc1 in injured fins *versus* control. Upon injury, not only is Smoc1 re-expressed in the fin, but also becomes present throughout the fin (Fig. 4C). Further, by quantifying Smoc1 intensity in fins across different timepoints, we observed a linear correlation between Smoc1 and the volume of the fin, highlighting that Smoc1 expression scales with fin size (Fig. 4D, Fig. S5C)^9,30^.

Next, we asked whether growth rate adaptation upon injury is dependent on these BMP-Smoc signaling dynamics. Our previous work showed that mutations in the *smoc1* gene lead to BMP signaling gradient scaling defects in the posterior side of the developing pectoral fin. Although only the posterior BMP gradient is perturbed, *Smoc1^-/-^* mutants show disrupted developmental growth, producing small fins^21^. To test whether we could perturb both BMP signaling gradients as the pectoral fin develops, we obtained *smoc2* mutants from the Zebrafish Mutation Project^67^. The *Smoc2^sa^*^24829^ allele generates a null mutation (Figs. S6A,E; Methods), and homozygous animals display similar BMP signaling gradients and Smoc1 expression to those of wildtype pectoral fins at 48 hpf (Fig. S6B-D). We then generated double homozygous *Smoc1^-/-^; Smoc2^-/-^*animals and asked, what fin growth consequences do the lack of Smoc1 and Smoc2 lead to, especially upon injury?

We first confirmed that *Smoc1^-/-^; Smoc2^-/-^*animals fail to develop fins with wildtype size, forming small fins (Fig. 4E-G), in line with known *Smoc1^-/-^* phenotypes^21^. Then by applying our injury assay to *Smoc1^-/-^; Smoc2^-/-^*pectoral fins and measuring their volume, area, and thickness over time, we unexpectedly observed that at 144 hpf, injured mutant fins surpass the size of uncut mutant controls, becoming of comparable sizes to wildtype fins (Fig. 4E-G, Fig. S7A-D). These phenotypes are restricted to injured fins, as *Smoc1^-/-^; Smoc2^-/-^* animals remain developmentally smaller than wildtype (Fig. 4I, Fig. S7E). By applying our mathematical model (Fig. 3H) to the *Smoc1^-/-^; Smoc2^-/-^* data, we find that uninjured mutant fins converge to a smaller area setpoint; in contrast, injured mutant fins acquire the size that is intrinsically set to wildtype (Fig. 4H), thus bypassing their genetic defects at the organ scale.

Last, to understand whether the observed growth compensation in injured *Smoc1^-/-^; Smoc2^-/-^* fins could be linked to altered BMP signaling, we performed comparative pSmad1/5/9 immunostainings across our datasets. Strikingly, by 12 hours post amputation (60 hpf), BMP signaling in injured *Smoc1^-/-^; Smoc2^-/-^* fins surpasses that of injured wildtype, persisting at 72 hpf (24 hpa), and declining 12 hours later, by 84 hpf (36 hpa). Like in injured wildtype fins, this upregulated BMP signaling in *Smoc1^-/-^; Smoc2^-/-^* mutants, is not only due to the presence of reacquired pSmad1/5/9 gradients, but also due to extraordinary signaling spatially localized abutting the amputation site. Injury thus induces additional BMP signaling that renders Smoc1/2 dispensable for fin growth control, restoring wildtype fin size despite existing genetic defects.

## Discussion

Our work reveals a role for injury-induced programs adapting organ developmental growth rates. An organ-wide increase in cell proliferation compensates for inflicted tissue loss, serving as a self-correction mechanism that can even override existing developmental defects. Thus, functional wildtype organ size and proportions are restored, while overall organismal growth is uninterrupted.

Previous studies reported differing accounts of how larval zebrafish pectoral fins recover from inflicted damage^51,68^. Here, using a precise UV-laser microdissection protocol, we establish the larval pectoral fin as a quantitative, reliable model for studying the molecular, cellular and tissue-level mechanisms that instruct organ growth adaptation. By reproducibly removing 30% of the fin at 48 hpf, we find consistent morphology recovery within four days (Figs. 1-2, 3B-D). Measurements from injuries that remove variable portions of the fin (Figs. 3E-F) can be concisely rationalized by a theoretical model implementing feedback control of growth (Fig. 3H, Supplementary Notes). From this, we propose that fin size is sensed and adjusted towards an organ-specific target area (Figs. 3I, S3E). Our model also clarifies that the adaptation of growth rates occurs on a timescale of several hours, recapitulating the experimental data.

Our data further shows that developing pectoral fins present signatures of embryonic robustness, where injury, and consequent cell loss, is compensated by increasing proliferation and extracellular space in an organ-wide manner (Figs. 2F-G, J-K). In contrast, injury triggers specific cellular responses that bypass developmentally active patterning. Strikingly, these responses restore wildtype fin size in mutants that usually cannot adapt their growth rates and develop small fins (Fig. 4). These results align with current reports in newts, where *Fgf10* mutants develop severely defective hindlimbs yet regenerate morphologically-resembling wildtype limbs after amputation. In that system, the blastema restores expression of *Fgf10*-regulated genes rather than reproducing the loss-of-function mutant developmental program^48^. Further, in unperturbed zebrafish larval finfolds, tissue regeneration enhancer elements (TREEs)^69,70^ appear to become engaged in response to genetically-encoded developmental defects, suggesting that regeneration-specific programs can be activated to protect against deleterious phenotypes^47^. Together, our findings extend these works and quantitatively argue that cells harness context-specific growth cues, adapting their growth rates to ensure functional organ shapes.

Beyond recovery of organ size to the adjusted developmental stage after injury, we observe also recovery of organ structure and shape, measured as tissue and organ proportions (Figs. 2G; 3D). We propose that injury-induced programs override organism-level developmental blueprints, and instead deploy an organ-level growth controller with an intrinsic setpoint corresponding to wildtype fin area (Figs. 3H-I; Fig. 4H). In injured *Smoc1^-/-^;Smoc2^-/-^*mutants, this mechanism is sufficient to bypass the developmentally encoded, but defective, BMP-Smoc feedback loop, triggering the injury controller that restores wildtype fin size. Intriguingly, recent work on adult zebrafish caudal fins shows that size memory is compromised in *longfin* mutants^40^. After amputation, these animals restore neither their own mutant pre-injury fin proportions nor wildtype ones. Integrated with our work, this suggests that injury signals override a defective patterning input (as in *Smoc1^-/-^;Smoc2^-/-^* mutants), but not a compromised growth effector (as in *longfin*). Deciphering the mechanisms that set intrinsic organ target size in development, while relating those to the cues instructing positional information and memory remains an exciting frontier to be explored.

Last, our findings position BMP signaling as a common effector of organ growth control, propagated by developmental gradient scaling^21,28^ or, independently, activated by injury. This dual access to a shared signaling effector may canalize organ size, ensuring the growth target is reached even in presence of perturbations or when the developmental route fails. Future work addressing the possible link between injury signals, canonical cellular stress responses (*e.g.* proteostasis), regenerative programs, and BMP signaling levels will clarify the molecular dynamics of cell proliferation and fin growth rate adaptation. Our study opens an avenue for resolving the spatiotemporal interplay in organ growth canalization routes driving developmental robustness.

## Acknowledgements

We thank Heewon Cho for initial efforts on performing Smoc1 immunostaining characterization in injured fins and Joseph Antiochus for help with initial genotyping of *Smoc2^sa^*^24829^ mutants. We are grateful to the following MPI-CBG and PoL facilities: Light Microscopy, Technology Development Studio, Cell Technologies, and Zebrafish Unit. This work was supported by the Max Planck Society (C.P., N.B., L.R., S.G., V.K., S.K., S.J., E.A., and R.M.) and the Deutsche Forschungsgemeinschaft (DFG, German Research Foundation) under Germany’s Excellence Strategy - EXC-2068-390729961 - Cluster of Excellence Physics of Life of TU Dresden (to all), a DFG Heisenberg grant #421143374 (B.M.F.), a DFG Individual Research Grant #548220137 (R.M.), and DRESDEN-concept support as part of the Excellence Strategy of the Federal and State Governments of Germany (R.M.). M.K. acknowledges support from the Studienstiftung des Deutschen Volkes.

## Author contributions

C.P., L.R., R.M., S.G., S.K., S.J., and V.K. performed experiments and analysed data. M.K., L.R., R.M., and S.G. developed analysis pipelines. B.M.F. and M.K. developed the theory and fin measurements computational pipeline. S.K. obtained qPCR samples. E.A., N.B. and S.K. performed cloning and established the *Tg(mylz2:QF2_myl7:GFP)^cbg36Tg^*and *Tg(QUAS:GFP_cryaa:CFP)^cbg35Tg^* zebrafish lines. B.M.F. and R.M. developed the project and wrote the manuscript. All authors read, edited and approved the final version of the manuscript.

## Data, code and materials availability

Code is available at: https://github.com/Coolix99/ZF_PF_Geometry. Plasmids and zebrafish lines generated in this study are available upon request to the lead contact, R.M.

## Declaration of generative AI and AI-assisted technologies in the writing process

During the preparation of this work, the authors used Claude 4.8 to edit the manuscript for spelling, grammar and clarity. The authors reviewed and edited the content as needed and take full responsibility for the content of the manuscript.

