## Supplementary Figures S1 to S7 for "Developmental growth rates adapt to enable self-correction of organ morphology after injury"

Shivani Gunnan, Maximilian Kotz, Lucas Ribas et al.

The PDF file includes:

- Supplementary Figures S1 to S7
- Materials and Methods
- Supplementary Theory Notes
- Supplementary Tables
- References of Supplementary Materials

Other Supplementary Materials for this manuscript include the following:

- Interactive 3D protein structure of wildtype Smoc1:BMP2a:BMP2a complex
- Interactive 3D protein structure of mutant Smoc1$^{-/-}$:BMP2a:BMP2a complex

### Supplementary Figures

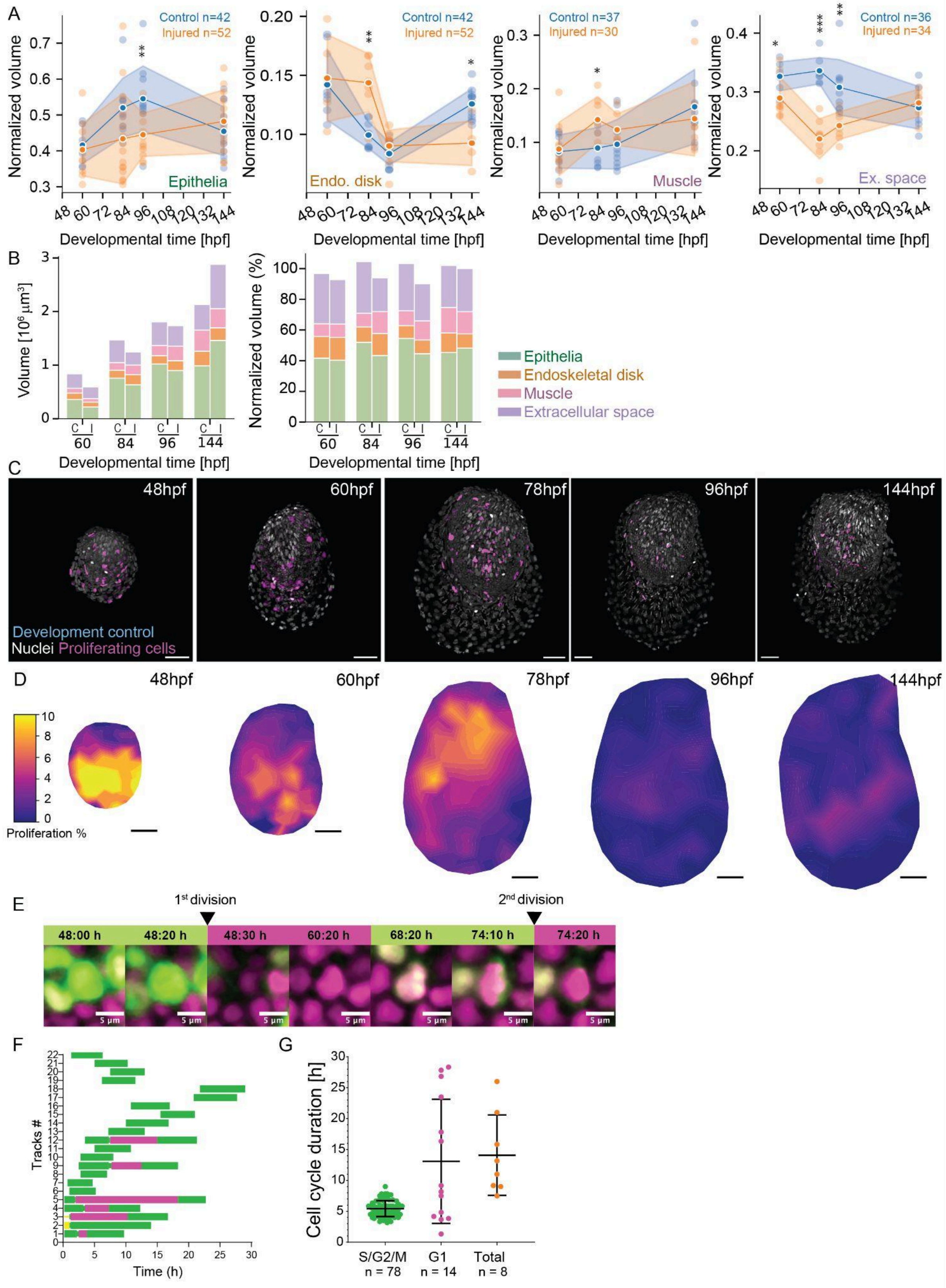


***Figure legend in next page***

**Figure S1. Tissue and cellular contributions to larval pectoral fin growth.**

**(A)** Tissue volume normalized to individual fin volumes comparing control (blue) versus injured (orange) fins for the epithelia, endoskeletal disk, muscle, and extracellular space, between 60-96 hpf. Note that during these stages the larval pectoral fin also contains a vascular loop (primitive pectoral artery[1]) and scattered fibroblast-like cells[2], whose volume contributions were not accessed, as these were deemed smaller than the other analysed fin components. Mean values connected by solid lines; shading indicates ±SD. All statistics: *$p \leq 0.05$, **$p \leq 0.01$, ***$p \leq 0.001$; two-tailed, unpaired, non-parametric Mann-Whitney tests. **(B)** *Right:* Volume of each pectoral fin tissue, in control (C) and injured (I) conditions, between 60-96 hpf. *Left:* Relative volume contribution (%) of each pectoral fin tissue normalized to individual fin volume, in control (C) and injured (I) conditions, between 60-96 hpf. Tissue color matches legend in Fig. 2A. Grey dash boxes indicate 100% mark. **(C)** Pectoral fins from transgenics labeling all nuclei (H2B-mCherry, white) and proliferating cells (mAG-zGeminin$^+$, magenta), during unperturbed development. **(D)** Spatial maps of proliferation in developing fins, from 48 hpf to 144 hpf. Color code shows the percentage of zGeminin$^+$ nuclei (indicative of a relative growth rate in %). zGeminin$^+$ nuclei were projected onto fin midsurface and averaged over $n$ = 8-17 fins per condition (Methods). Scale bars: 50μm. **(E)** Zoom-in sequence images of a single cell nuclei (center of frame) tracked from light sheet long-term live imaging, highlighting two cell divisions. Green: mAG-zGeminin signal. Magenta: H2A-mCherry signal. **(F)** Representative time course of 22 cells tracked from a 30h dataset displaying the cell cycle duration of S/G2/M (green, from mAG-zGeminin transgenics) and G1 phases (magenta, by exclusion of mAG-zGeminin signal). **(G)** Comparison between duration of cell cycle phases for cells of developing pectoral fin. Mean ± SD. n, number of cells. For all images: anterior, left; distal down.

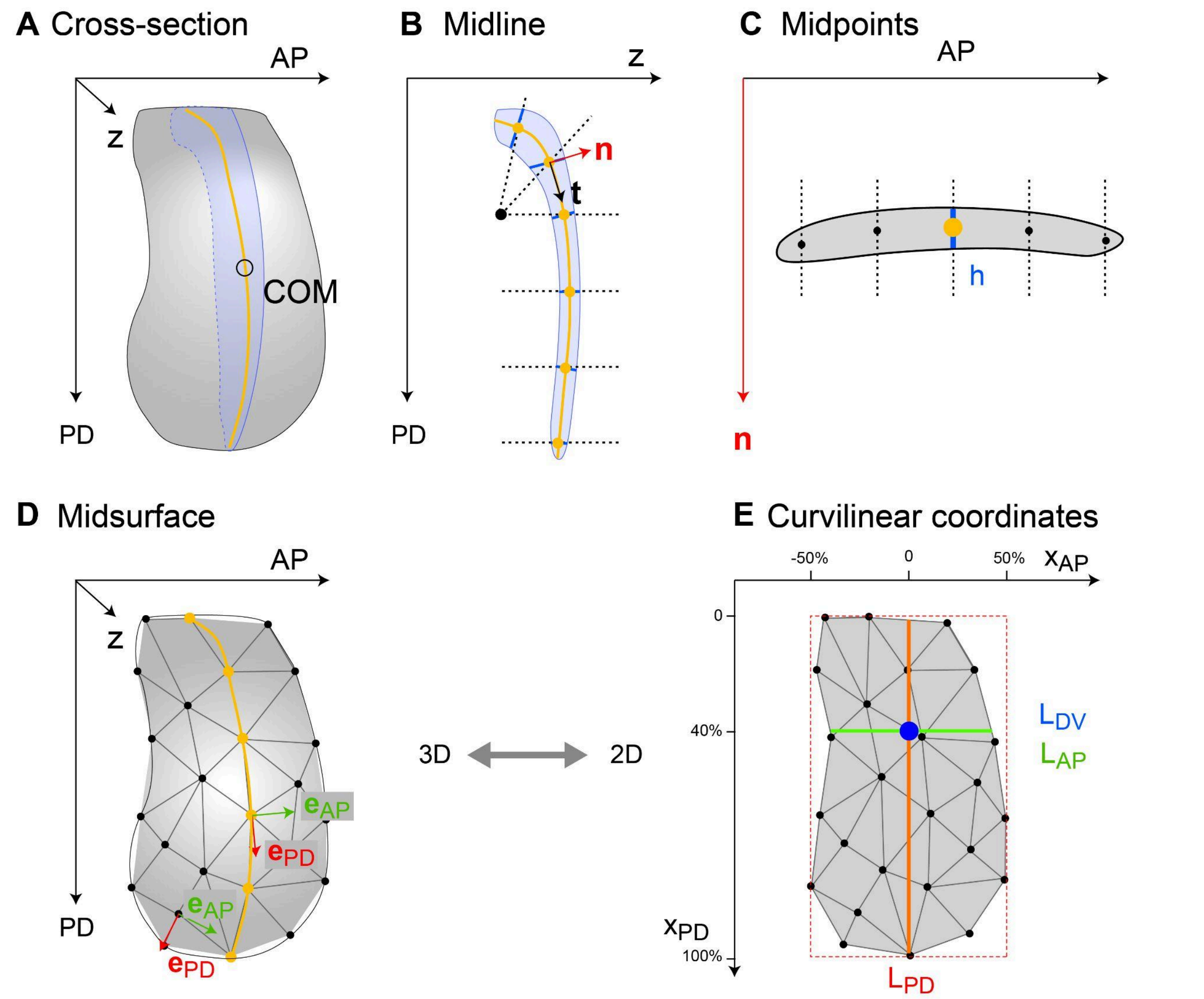


**Figure S2. Computation of fin midsurface and axis coordinates.**

**(A)** Cross-section (blue) of binary fin mask (gray) passing through its center-of-mass (COM) and normal to a manually annotated anterior-posterior axis (AP). **(B)** Midline (orange) of blue cross-section from A, determined from midpoints of line segments given by intersections of rays (black dotted) starting from a manually selected reference point (black) and the fin mask. The midline defines a local tangent vector **t** and a local normal vector **n**. **(C)** Orthogonal cross-section passing through a point of the midline (orange dot) and normal to **t**. The intersections of lines parallel to **n** (dotted lines) with this cross-section define line segments. The midpoints of these segments define a set of points in the center of the cross-section (black dots), which will be used to compute the fin midsurface. **(D)** Fitting a surface to the midpoints from all orthogonal cross-sections, as shown in C, defines the midsurface of the fin, represented as a triangulated surface in 3D. The midline (orange) and local unit vectors $\mathbf{e}_{PD}$ and $\mathbf{e}_{AP}$ are highlighted. **(E)** Curvilinear coordinates of midsurface $x_{PD}$ and $x_{AP}$ (obtained by integrating the vector $\mathbf{e}_{PD}$ along the midline to define $x_{PD}$, and integrating $\mathbf{e}_{AP}$ starting from the midline to define $x_{AP}$, see C). This 2D-cartographic projection of the fin midsurface allows for defining linear lengths: $L_{AP}$: anterior-posterior length measured at 40% of proximal-distal relative position, computed as extension $L_{AP,\ max}$: longest length along the $x_{AP}$-coordinate, $L_{AP,40\%}$: anterior-posterior length along the $x_{AP}$-coordinate using the line segment passing through a reference point (blue, defined below), $L_{PD}$: proximal-distal length along the $x_{PD}$-coordinate using the line segment passing through the reference point, $L_{DV}$: dorsal-ventral length, computed in the 3D fin mask from a line segment passing through the reference point (mapped back from the flattened midsurface to the curved midsurface) and parallel to the local normal vector **n**. The reference point was chosen at a relative position ($x_{AP}$ = 0, $x_{PD}$ = 40%) within a rectangular bounding box with edges parallel to the $x_{AP}$- and $x_{PD}$-coordinate axes and dimensions $L_{AP,\ BB}$ and $L_{PD,\ BB}$, where $x_{AP} = -L_{AP,\ BB}/2$, $x_{AP} = +L_{AP,\ BB}/2$, $x_{PD} = 0$, and $x_{PD} = -L_{PD,\ BB}$ specify the most anterior, most posterior, most proximal, and most distal edge of this bounding box, respectively.

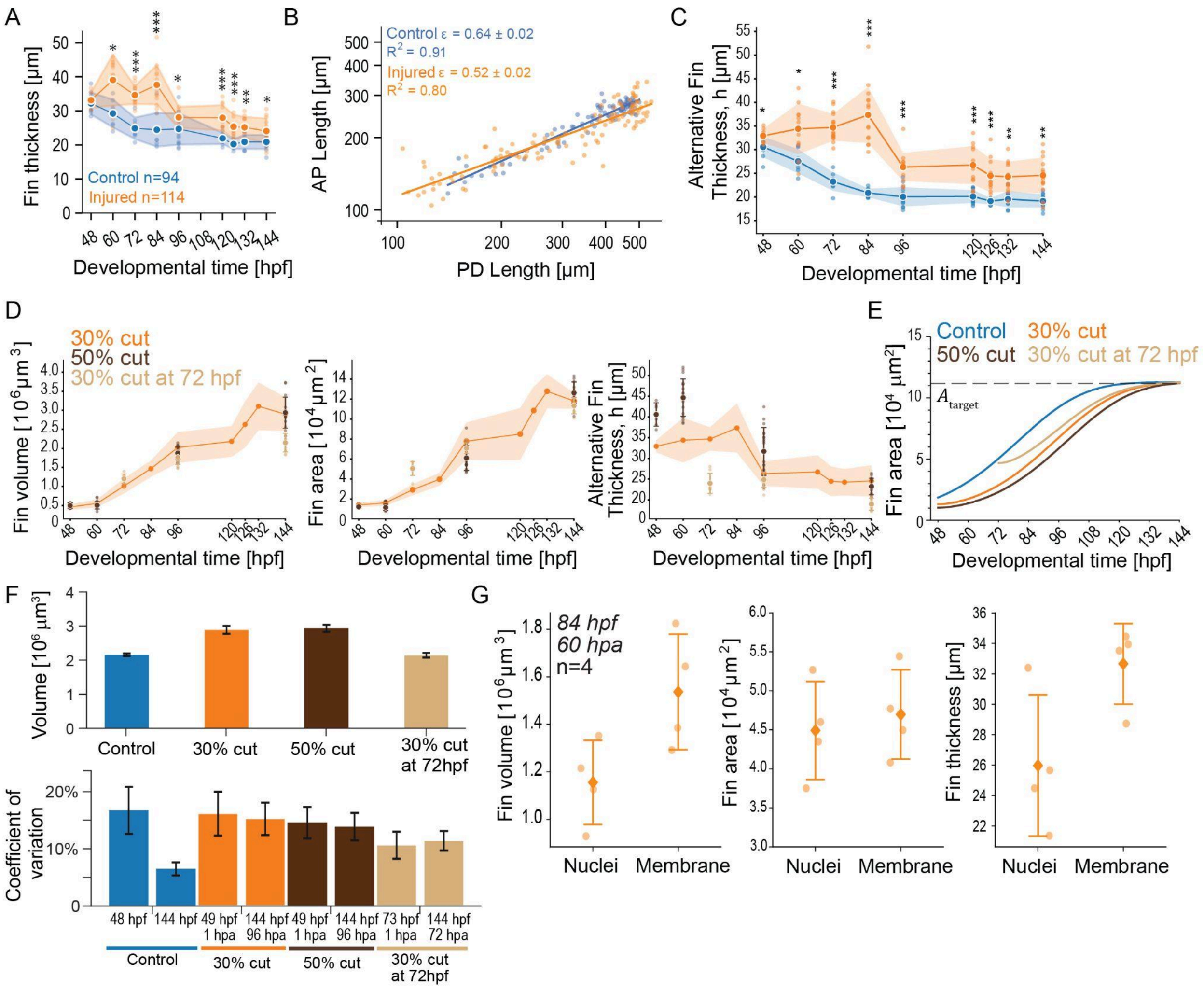


**Figure S3. Injury decouples fin growth across developmental axes.**

**(A)** Fin thickness, computed by dividing the measured fin volume (Fig. 1F) by fin area (Fig. 3C), as a function of time, in injured (orange) or control (blue) fins. **(B)** Log-log plot of anterior-posterior (AP) versus proximal-distal (PD) lengths to determine the fin anisotropic growth rate, ε (slope value) ± SEM, in injured (orange) and control (blue) fins. Line, powerlaw fit with goodness of fit ($R^2$). **(C)** Alternative fin thickness, h, in injured (orange) or control (blue) fins. See Methods. **(D)** From *left to right*, fin volume and thickness as a function of time, in the tested three injury conditions: 30% cut at 48 hpf (orange, from panel C, and Figs. 1G, 3C), 50% cut at 48 hpf (brown), and 30% cut at 72 hpf (beige). Mean ± SD shown. Number of fins analysed: n=114 (30% cut at 48hpf), n=40 (50% cut) and n=32 (30% cut at 72 hpf); 6-15 per condition. **(E)** Model fits (from Fig. 3I) for fin area growth across conditions superimposed. Note how the fin's target area is always reached, irrespective of the perturbation. **(F)** *Top:* Comparison of fin volume at 144 hpf for control (blue) and the three tested injury scenarios. Mean ± SD shown. *Bottom:* Coefficient-of-variation (i.e. sample standard deviation normalized by sample mean at

48 and 144 hpf) for control (blue) fins and the three tested injury conditions. Mean ± SEM. Number of fins analysed: n=23 (uncut control), n=22 (30% cut at 48hpf), n=15 (50% cut), n=20 (30% cut at 72 hpf); 6-15 per condition. **(G)** From *left to right*, comparison of fin volume, area, and thickness from 3D fin masks obtained from transgenic fish labelling nuclei (H2B-mCherry signal) and membranes (claudinb:Lyn-GFP signal) simultaneously, in injured fins at 84 hpf (60 hpa). Note that different labels render different volumes and consequently thicknesses, but fin area is comparable across the two signals. Mean ± SD shown. All statistics: **p*≤0.05, ***p*≤0.01, ****p*≤0.001; two-tailed, unpaired, non-parametric Mann-Whitney tests.

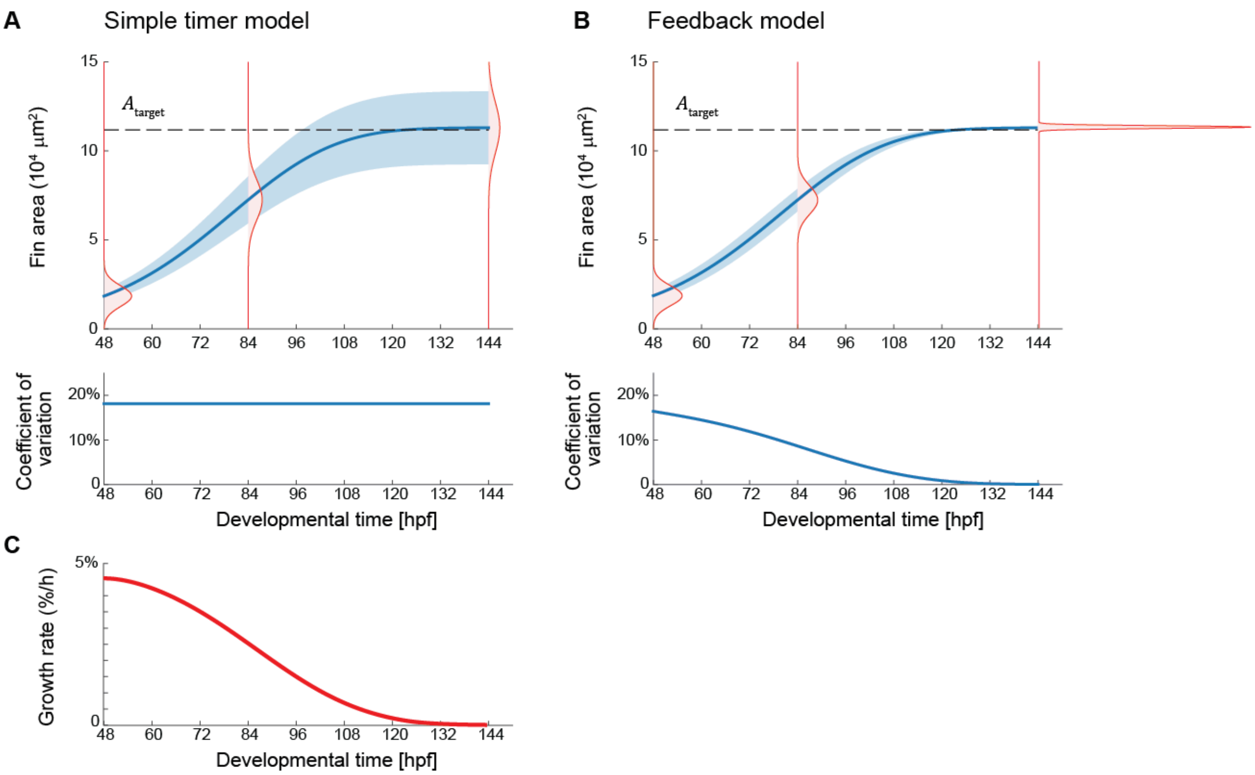


**Figure S4. Predicted change in size variability for two growth control models.**

**(A)** In a simple timer model, area $A(t)$ grows according to a prescribed growth rate, without dynamic adaptation. For this model, the coefficient-of-variation CV (i.e., standard deviation normalized by mean) stays constant in time. In the upper panel, the computed mean area $A(t)$ (mean: solid blue, ±SD: blue shade) is shown, together with distributions of fin area $p(A,t)$ at selected time points $t$ (red). In the lower panel, the corresponding coefficient-of-variation CV($t$) is shown. The growth rate was chosen to reproduce the same growth dynamics as shown for the fit for control condition in Fig. 3I, see panel C. **(B)** In a feedback-control model, area $A(t)$ grows according to an adaptive growth rate that dynamically changes in response to the ratio between the current area $A(t)$ and a target area $A_{target}(t)$ that serves as a setpoint, see Eqs. (1-3) (Supplementary Theory Notes). This feedback-control model reproduces exactly the same mean dynamics as the simple timer model shown in panel A (with same mean growth rate as used in panel B, shown in panel C), yet results in decrease of the coefficient-of-variation CV with time. **(C)** Relative growth rate $g(t)$ corresponding to the fit shown in Fig. 3I. Parameters: $A(t_0)$ = $A_0$ = 1.8·$10^4$ μm$^2$ (Fig. 3I), CV($t_0$) = 18% (Fig. 3G) at $t_0$ = 48 hpf.

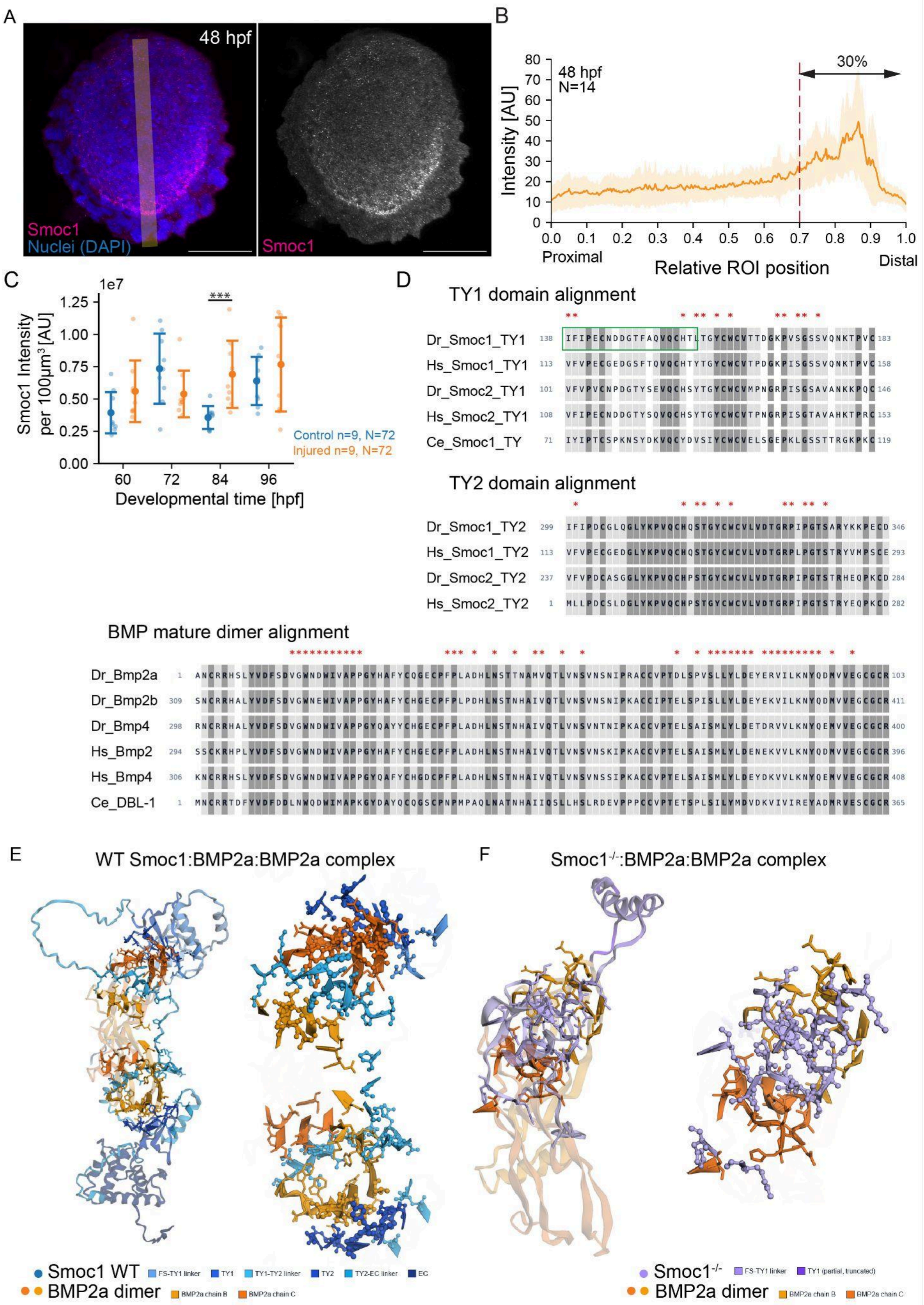


***Figure legend on next page***

**Figure S5.** ***In silico*** **modeling shows direct interactions between zebrafish Smoc1 and BMP, lost in the** ***Smoc1$^{ug104}$*** **mutant.**

**(A)** Anti-Smoc1 (magenta) immunostaining in the pectoral fin at 48hpf. The region of interest (ROI) along the proximal-distal fin axis is shown in yellow. Anterior, left; distal down. Scale bars: 50µm. **(B)** Average Smoc1 intensity along the ROI defined in A versus relative position (normalized ROI length). Shading, SD. Red dash indicates region where UV-laser microdissection is applied to the fins, removing 30% distal-most part of this organ, encompassing a large portion of the Smoc1 expression. **(C)** Average Smoc1 intensity per 100µm$^3$ in control (blue) or injured (orange) pectoral fins, at different developmental times. Mean ±SD. Statistics: ***$p \leq 0.001$; two-tailed, unpaired, nonparametric Mann-Whitney tests. N, total number of fins analysed; n, number of fins per timepoint. **(D)** Multiple sequence alignment of the TY domains of Smoc1 homologs using pairwise BLOSUM62. Dark grey boxes indicate identical residues, light grey boxes indicate conserved residues among homologs. Red asterisks (*) mark the residues at the interface between Smoc1 and BMP, as identified via ColabFold (see below and [3]). Green box indicates the zebrafish *Smoc1$^{ug104}$* mutant allele[4], which leads to a frameshift from aa134, generating a truncated protein. **(E)** Predicted structures of complexes formed between the wildtype secreted form of DrSmoc1 (blue) and a mature DrBMP2a dimer (yellow, orange). *Left*, full complex shown; *Right*, only contact residues (≤ 5 Å) shown. Predictions are similar between DrSmoc1 and DrSmoc2, but only DrSmoc1 is shown here. Predictions are also similar between DrBMP2b and DrBMP4, but only DrBMP2a is shown here. Note that each of the BMP ligands is predicted to make a similar interaction with a TY domain of its corresponding Smoc1 homolog. **(F)** Predicted structures of complexes formed between the mutant secreted form of DrSmoc1$^{-/-}$ (purple) and the mature DrBMP2a dimer (yellow, orange). Note that in Smoc1$^{-/-}$ mutants, none of the wildtype interactions are preserved. However, the resulting Smoc1 protein (if not degraded) can still bind to one of the BMP molecules, at positions not conserved in wildtype. *Left*, full complex shown; *Right*, only contact residues (≤ 5 Å) shown. BMP, bone morphogenetic protein; EC, extracellular calcium-binding domain; FS, follistatin-like domain; Smoc, secreted modular calcium-binding protein; TY, thyroglobulin type-1 domain.

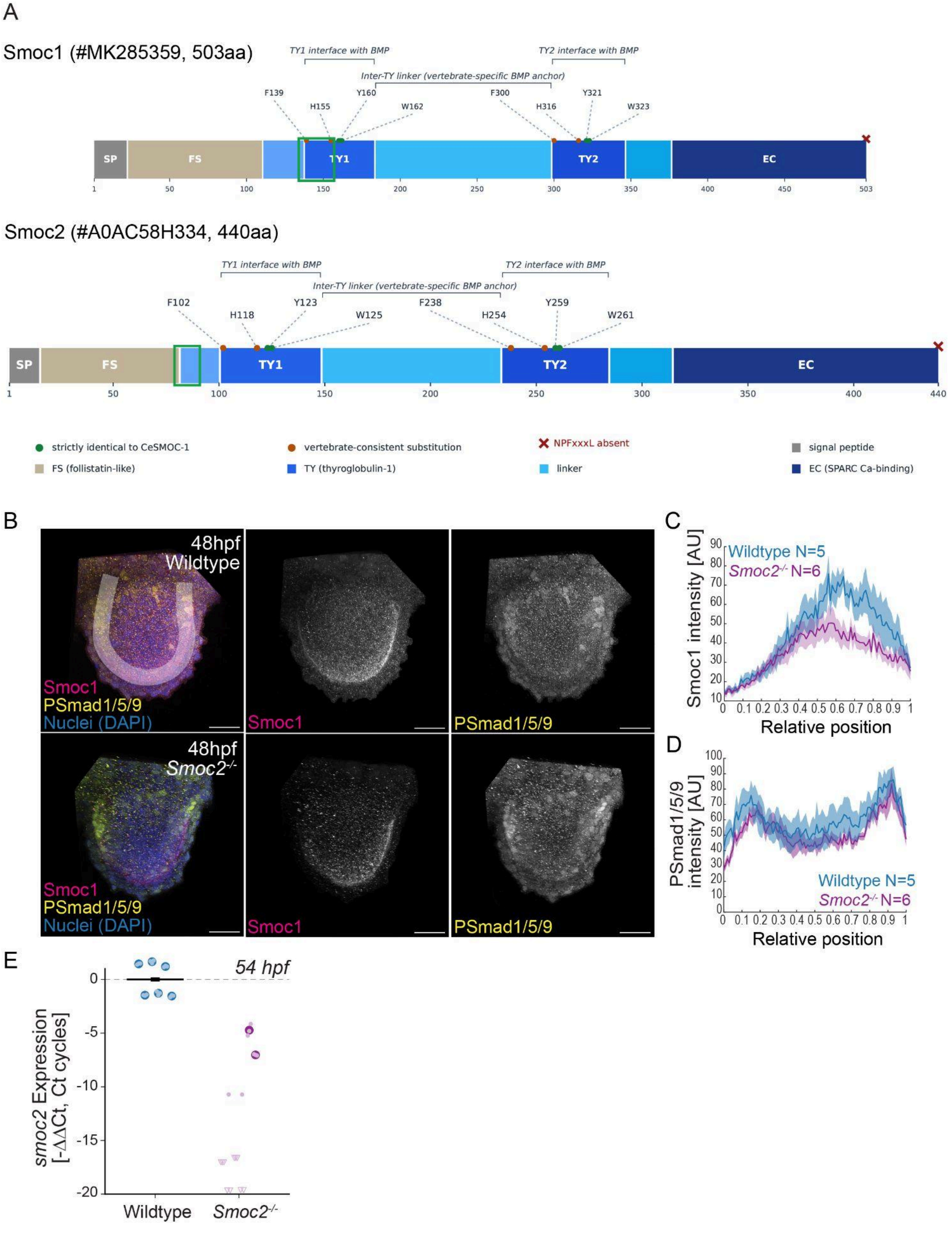


***Figure legend in next page***

**Figure S6. Characterization of *Smoc2$^{sa24829}$* mutants.**

**(A)** *Top:* DrSmoc1 protein structure. Green box indicates the zebrafish *Smoc1$^{ug104}$* mutant allele[4], which leads to a frameshift from aa134, generating a truncated protein. *Bottom:* DrSmoc2 protein structure. Green box indicates the zebrafish *Smoc2$^{sa24829}$* mutant allele, which leads to intron-1 being retained, leading to a premature stop at aa92. Note that in zebrafish Smoc1 TY domains, Y160, W162, Y321, W323 are strictly identical to the *C.elegans* Smoc1, shown to bind BMP[3]. The zebrafish Smoc1 is also similar to the Human sequence where a NPFxxxL motif is absent from the EC C-terminus (red cross). Instead, two F-x-x-x-L submotifs at F220/F222 sit in the TY1-TY2 linker, precisely the region that AlphaFold predicts as the vertebrate-specific BMP anchor in HsSMOC1, as shown in [3]. **(B)** Anti-Smoc1 (magenta) and anti-Phospho Smad 1/5/9 (yellow) immunostainings in pectoral fins from wildtype (*top*) and *Smoc2$^{sa24829}$* homozygous mutants (*bottom,* Smoc2$^{-/-}$) at 48 hpf. Anterior, left; distal down. Scale bars: 50µm. **(C-D)** Average intensity of normalized Smoc1 (C) or Phospho-Smad 1/5/9 (D) immunostaining signal versus relative position (ROI midline, white) in wildtype (blue) versus *Smoc2$^{sa24829}$* homozygous mutants (purple). Shading, SD. **(E)** qPCR determination of *smoc2* expression levels in wildtype (blue) versus *Smoc2$^{sa24829}$* homozygous mutants (purple), at 54 hpf. Large dots, biological replicate mean; small dots, technical replicates; triangles, non-detected expression at Cq 45; grey dash, wildtype mean reference. Note that statistical comparisons were not made (or fold change calculation) due to inconsistent *smoc2* detection in *Smoc2$^{-/-}$* biological replicates, indicative of a null mutation leading to degradation of respective mRNA in those samples. *N*, total number of fins analyzed.

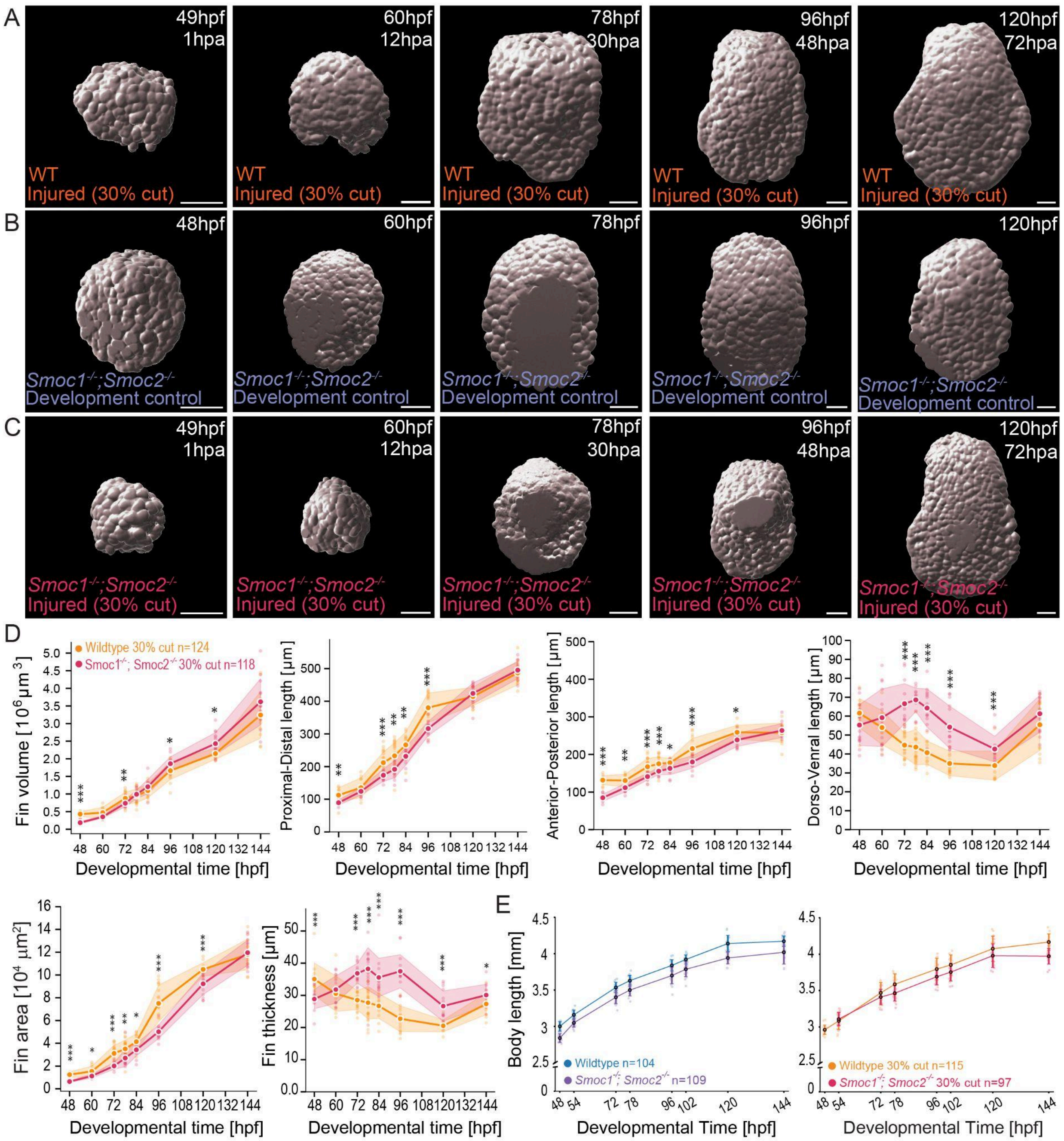


**Figure S7. Pectoral fin and body growth dynamics in *Smoc1$^{-/-}$;Smoc2$^{-/-}$* mutants.**

**(A-C)** 3D surface masks of pectoral fins live imaged at different timepoints in 30% cut wildtype fins (A), *Smoc1$^{-/-}$;Smoc2$^{-/-}$* developing fins (B), and *Smoc1$^{-/-}$;Smoc2$^{-/-}$* injured fins (C). Note that masks were obtained from nuclei signal (H2B-mCherry). For all images: anterior, left; distal down. Scale bars: 50µm. **(D)** *Top row, from left to right:* pectoral fin volume, fin lengths along proximal-distal, anterior-posterior or dorso-ventral fin axes as a function of time, in injured wildtype (orange) *versus Smoc1$^{-/-}$;Smoc2$^{-/-}$* mutants (pink), after after 30% cuts. *Bottom row,* *from left to right:* pectoral fin area and thickness in injured wildtype (orange) *versus* *Smoc1*$^{-/-}$*;Smoc2*$^{-/-}$ mutants (pink), after 30% cuts. **(E)** Larval body length as a function of time in developing wildtype and *Smoc1*$^{-/-}$*;Smoc2*$^{-/-}$ mutants, with uncut (*left:* WT, blue; mutant, purple) or injured (*right:* WT, orange; mutant, pink) pectoral fins. Mean values connected by solid lines. For all plots: bars or shading indicate ±SD. *n*, number of fins (D) or larvae (E); 8-24 per timepoint. All statistics: **p*≤0.05, ***p*≤0.01, ****p*≤0.001; two-tailed, unpaired, non-parametric Mann-Whitney tests.

### Materials and Methods

##### Ethics Statement

This study followed European Union directives (2010/63/EU) and German law, with license #TVV52/2021 - ‘Generierung von Zebrafischlinien zur Untersuchung der Größe und Form von Organen und Organellen‘. Genetic engineering work was carried out in an S1 area (MPICBG, S1-Labore 4., Az.: 54-8451/103, project leader Rita Mateus), following guidelines according to Section 21, Paragraph 1 of the German Genetic Engineering Act, and within projects 01-03 from the Mateus laboratory, according to Section 28 of the German Genetic Engineering Safety Ordinance (GenTSV).

##### Zebrafish lines and maintenance

All zebrafish (*Danio rerio*) lines were maintained in a recirculating system with a 14 h/day, 10h/night cycle at 28°C. Crosses were performed with 3- to 12-month-old adults. Embryos were kept in E3 zebrafish embryo medium[5] at 28.5°C until reaching the desired developmental stage. We used the following lines in WT AB background: Tg(EF1a:mAG-zGem(1/100))$^{rw0410h}$ [6], referred to as mAG-zGeminin; Tg(Xla.Eef1a1:hist2-mCherry)[7], referred as H2B-mCherry; Tg(BmpRE:EGFP)$^{pt509}$ [8], referred as BRE:GFP; Tg(-8.0cldnb:lynGFP)$^{zf106}$ [9], referred as claudinb:Lyn-GFP; Tg(actb2:LY-tdTomato)$^{zf2254Tg}$ [10], referred as b-act:lyn-tdTomato; *Smoc1*$^{ug104}$ [4]; Tg(-7.2sox10:MA-RFP)$^{ya300Tg}$ [11]; and Tg(ubb:SEC-Rno.Ncan-EGFP)$^{uq25bhTg}$ [12].

##### *Smoc2*$^{sa24829}$ tilling mutants

The mutant line *Smoc2*$^{sa24829}$ was generated by the Zebrafish Mutation Project[13]. The *Smoc2*$^{sa24829}$ allele is an essential splice site mutation affecting the donor site of exon 1 of *smoc2* (chr13:43,263,999 T>G; GRCz10) (ENSEMBL ID: ENSDARG00000088157). In addition to the *Smoc2*$^{sa24829}$ mutation, 28 other mutations had been mapped to the F1 male fish from which F2 *Smoc2*$^{sa24829}$ animals were generated. Hence, prior to usage of the line, extensive genotyping was performed on adult F2 animals for these other mutations, enabling identification of the animal with the least amount of additional mutations to *Smoc2*$^{sa24829}$. Subsequently, line propagation carrying only the *Smoc2*$^{sa24829}$ mutation was performed by outcrossing the identified *Smoc2*$^{sa24829}$ mutants with wildtype AB. F4 generation *Smoc2*$^{sa24829}$ heterozygous animals were used for crosses with homozygous *Smoc1*$^{ug104}$, and double mutant homozygous animals were subsequently generated by incrossing.

##### Transgenic line generation

Transgenesis was performed using the Tol2 transposon system and Gateway cloning[14–16] technology. For the *Tg(mylz2:QF2_myl7:GFP)*$^{cbg36Tg}$ line, a new 5’ entry clone with the *mylz2* promoter sequence was generated. The full length 1943bp *mylz2* promoter sequence[17] was amplified using the following primers: FWD 5’-GGGGACAACTTTGTATAGAAAAGTTGTAATTCGCCACAGAGGAATGAGC-3’; REV 5’-GGGGACTGCTTTTTTGTACAAACTTGTTGTACTTGAGGGGCTTATATACTGACGA-3’, using wildtype AB genomic DNA, introducing recombination sites. The PCR product was then purified using a PCR clean up kit (Promega #A9281) and recombined with pDONOR P4-P1r (Tol2kit #213) in a BP reaction (BP Clonase II enzyme mix, Invitrogen, #11789020). The resulting 5’ entry clone was was then recombined in a LR reaction (LR Clonase II Plus enzyme, Invitrogen, #12538120) using pDEST-Tol2pA2_myl7:EGFP (R4-R3) (Tol2kit #395), pME-QF2 (Addgene #83307) and p3E-polyA (Tol2kit #302) to generate the final construct T2_mylz2:QF2_pA_myl7:GFP. For the *Tg(QUAS:GFP, cryaa:CFP)$^{cbg35Tg}$* line, the final construct QUAS:GFP_pA_betacrystallin:CFP was a kind gift from Marnie Halpern (Addgene #81181). One-cell stage WT AB embryos were injected with 25 pg of each of the final constructs, 25 pg of Tol2 transposase mRNA, and 10% phenol red. At 2 days postfertilization, injected embryos were screened for GFP/CFP expression in the heart (driven by the *myl7* selection marker) or lens (driven by the *cryaa* selection marker) using an Olympus SZX16 fluorescence microscope and subsequently grown to adulthood. Founder fish were identified by screening F1 progeny for selection marker fluorescence, resulting from outcrosses with WT AB fish. F2 or later generations in WT AB background were used for this study's data acquisition.

**mRNA synthesis**
For Tol2 transposase mRNA synthesis, the pCS2FA-transposase plasmid[14] was linearized with NotI (New England Biolabs) and transcribed using the SP6 mMessage Machine Kit (Invitrogen, #AM1340) according to the manufacturer's protocol.

**Genotyping**
Genomic DNA from individual larvae or clipped tail fins from adult zebrafish were extracted using the Kapa Express Extract kit (Kapa Biosystems) according to the manufacturer's protocol. This was followed by performing PCR with KAPA2G Robust HotStart ReadyMix (Kapa Biosystems). For *Smoc1$^{ug104}$* genotyping, the following primers were used: FWD 5'-CAAGAATCCATTCCCCCC-3'; REV 5'-CTTTAGGATTCGTTGATGTCAAAAG-3'. For *Smoc2$^{sa24829}$* genotyping, the following primers were used: FWD 5'-CTGGAGCTCCTCGAAAGC-3'; REV 5'-TGTGATTTTTAAATTGCTTTGATTTG-3'. Resulting PCR products from either mutant were sent for sequencing and sequences analyzed using Snapgene (v7.1).

**Cellmask treatments**
Double transgenic QUAS:GFP; mylz2:QF2 larvae (labelling the muscle) were incubated for 1 hour with the lipophilic dye Cellmask DeepRed plasma membrane stain (Invitrogen #C10046), diluted at a 1:1000 in E3 embryo medium, prior to live imaging.

**UV-laser pectoral fin microdissection**
Pectoral fin amputations were performed using a Zeiss Axio Observer.Z1 inverted microscope with a focused UV-laser (355nm), a PALM MicroBeam System, and a AxioCam Icc1 CCD camera. A 2.5x/0.08 NA EC Plan-Neofluor (Zeiss) air objective was used to locate samples and record positions in the Navigation Window mode of the software. Laser cutting was performed using a 20x/0.4 NA LD Plan-Neofluar (Zeiss) air objective. The following settings were used for UV-Laser cutting: Speed - Cutting: 130; Laser – Cut Energy: 57; LPC Energy: 57; Focus (1): 63; Focus (2) – 70; Cycles: 1. Previous to microdissection, larvae were anaesthetized in 0.1% MS-222 (Sigma) diluted in E3 medium and mounted in 35mm glass-bottom dishes (MatTek). Note that larvae were manually aligned such that their right-sided pectoral fins were against the

glass bottom, facing the objective. Brightfield illumination using the 2.5x objective was used to locate the fish and their positions were recorded in the Navigation Window. Once the pectoral fin was located, the 20x objective was used for focusing, coupled with fluorescent excitation (470nm or 625nm) for precise transgene visualization. The measuring tool was used to draw a line along the proximal-distal (PD) axis of the fin, determining the position for amputation at 30% from the distal fin tip. Then, using the line cut tool, a parallel line was drawn perpendicular to the PD line (along the fin's anterior-posterior axis), setting the amputation position. The UV-laser beam was activated by pressing the cut bottom. In case amputation was unsuccessful, the focus was adjusted as necessary and the cutting was repeated. To limit excessive damage to the fin, five attempts per larvae were performed. If amputation would still be unsuccessful, the sample was discarded. Amputated larvae were transferred to fresh E3 medium and kept at 28.5 ºC until the desired developmental stage. The same protocol was followed for amputations at 50% from the distal fin tip at 48 hpf, and 30% from the distal fin tip at 72 hpf.

**Confocal live imaging**

Confocal live imaging was performed in embryos anaesthetized in 0.1% MS-222 (Sigma) diluted in E3 medium, using a Fluorodish (World Precision Instruments) and imaged in a Zeiss LSM880 AiryScan point scanning confocal microscope, equipped with a C-Apochromat 40x/1.2 water immersion objective.

**Light sheet live imaging**

Long-term live imaging was performed using a LS1 Live (Viventis Microscopy) light sheet inverted microscope with a Nikon Apo 25x/NA 1.1 water-immersion objective (with 0.75x magnification factor), running Viventis Microscope Control 2.0.0.5 software. Double transgenic mAG-zGeminin; H2A-mCherry larvae at 48hpf were anaesthetized in 0.1% MS-222 (Sigma) prior to imaging. Samples were mounted in a Viventis sample holder filled with E3 medium and 0.1% MS-222. The sample holder was immersed in a chamber containing water at 28.5ºC, with temperature controlled by a thermostat. Imaging of the pectoral fin was acquired for 150 z-stacks using a 1µm step size, every 10min for 30h (between 48 and 78 hpf) using the single-sided illumination mode. The laser beam had a thickness of 2.2µm and was aligned using the Lightsheet alignment advanced module from the software. The illumination intensity was of 100% combined with an OD1 illumination filter, giving an effective laser power of 10%. Images were acquired with an exposure time of 100ms for a 440µm range.

**Immunofluorescence**

Whole-mount immunostainings were performed as previously described[18]. The primary antibodies used were: mouse anti-Smoc1 (1:200, Abnova #H00064093-M03), rat anti-GFP (1:100, Santa Cruz Biotechnology, catalog ref# sc-101536.) and rabbit anti-PSmad1/5/9 (1:100, Cell Signaling ref#13820). The secondary antibodies used were: goat anti-mouse Alexa 594, goat anti-rat Alexa 488, and goat anti-rabbit 647 (all 1:500, Molecular probes). DAPI (#D9564, Sigma) was incubated after secondary antibody incubation at 1:1000 dilution. Immunostainings were repeated at least 3 times with different biological replicates, per marker and condition. Embryos were mounted for imaging in 80% Glycerol, 2% DABCO (Sigma) diluted in PBS. Images were acquired with an inverted Zeiss LSM880 AiryScan point scanning confocal microscope equipped with a C-Apochromat 40x/1.2 water immersion objective (Zeiss) or a Zeiss Celldiscoverer 7 automated imaging system equipped with a Plan-Apochromat 50x/1.2 water-immersion objective (Zeiss). For the latter, embryos were mounted in 96-well cell culture plates (Greiner Bio-One, Cat. No. 655090). Multi-track acquisition mode was used to minimize spectral overlap between fluorophores where appropriate. Z-stacks spanning the entire pectoral fin were acquired for all samples. Smaller fins were imaged as a single field of view, whereas larger fins were acquired using multi-tiled imaging with 10% overlap, and the resulting tiles were stitched to reconstruct the complete fin. Image acquisition was performed using ZEN Black 2.3 SP1 for the LSM800 system and ZEN Blue 3.6 (Carl Zeiss) for the Celldiscoverer 7.

**Total RNA isolation and quantitative real-time PCR (qPCR)**

For all gene expression analyses, tissue from 25 larvae was harvested per experiment and pooled for each biological replicate. All samples were analyzed in biological and technical triplicate for each gene. Total RNA was extracted from samples using Trizol reagent (Merck #93289) according to the manufacturer's protocol. cDNA was synthesized from 1 µg total RNA using the Superscript IV First Strand Synthesis System with ezDNAase kit (Invitrogen #18091150), following the manufacturer's protocol, using random Hexamers. qPCR was performed using a Roche LightCycler 96 and FastStart Essential DNA Green Master Mix. Cyclic conditions were: 10min at 95°C followed by 45 amplification cycles, each consisting of 10 s at 95°C, 10 s at 60°C, and 15 s at 72°C. Gene expression values were normalized using the elongation factor 1α (*ef1α*, NM_131263; *eef1a1l1* – Zebrafish Information Network) housekeeping gene and respective expression analysis were calculated using the 2^(−ΔΔCT) method[19]. Results were plotted using a custom Python script, adapting the superplot convention[20]. Two-tailed, non-parametric paired Welch tests were performed between the several conditions and controls. Primer sequences are listed in supplementary material Table S1.

#### QUANTIFICATION AND STATISTICAL ANALYSIS

**Body size quantification**

Body size was quantified from brightfield images of anesthetized larvae from 48 hpf to 144 hpf, across different datasets. Brightfield images were acquired with a Leica M165C stereoscope (zoom 2) and then processed using Fiji[21], where the body length of each individual larva was measured from head to distal tip of the caudal fin fold. Plotting, linear regression fitting, and statistical analyses between body lengths from different conditions along time were done using a custom made Python script. Statistical comparisons were performed using unequal-variance t-test, unpaired, two-tailed, pairwise Welch's *t*-tests between the different conditions. Statistical experimental details can be found in figure legends.

**Fin volume quantification**

Fin volume, $V$, was quantified using binary 3D masks obtained from nuclei (H2A-mCherry) or membranes (claudin:Lyn-GFP or cellmask-treated) confocal z-stacked images. The first step was to create 3D surfaces in IMARIS using a surface detail of 0.5 - 0.8 µm, a manual intensity threshold to cover the entire surface of the fin and by splitting touching objects with a seed points diameter of 2-3.5 μm. Next, regions of the body outside of the pectoral fin were manually deleted from this surface using the Edit/Selection/Delete tab in IMARIS. This allowed us to obtain a 3D surface that exclusively covers the fin. This surface was used to generate a 3D mask of the fin (all voxels inside the closed surface were retained, while all voxels outside were assigned the value 0), which was saved as a separate channel in the IMARIS generated image file. As the 3D mask produced by IMARIS contains voids between the nuclei signal, further processing to generate a smooth, continuous mask, without gaps, was necessary. For this, the mask was thresholded to create a binary mask of voxels with intensity greater than 1. Then a morphological closing operation was applied in each XY slice to smooth the mask and connect nearby regions. The closing operation consisted of a dilation operation followed by an erosion operation with a disc radius of 18 μm as the structural element. In this way, small gaps in the mask smaller than 1000 area units within each slice were also filled. Importantly, by doing this, the smaller disconnected regions were removed and only the largest connected component was retained thus returning a clean boolean mask of the fin. The final 3D masks were then manually verified and corrected by hand if necessary. Fin volume in $\mu m^3$ is quantified by multiplying the number of voxels inside the binary mask by the volume of voxels in $\mu m^3$. A typical voxel size in x, y, z-direction was 0.2μm, 0.2μm, 0.7μm, respectively. The same analysis pipeline was applied to fins with cell membranes labeled. For fins treated with cellmask, the threshold in IMARIS parameters was lower since the signal was much brighter. Statistical analysis and comparison between fin volumes was performed using either GraphPad Prism or Python using the statannot package[22] by carrying out non-parametric, unpaired, two-tailed Mann Whitney *t*-tests between the different conditions. Statistical experimental details can be found in figure legends.

**Pectoral fin compartment 3D segmentation and volume quantification**

Each of the constituent tissues/compartments of the pectoral fin were labelled with a specific transgenic reporter (Epithelia: *claudinb*:Lyn-GFP; Endoskeletal disk: *sox10*:MA-RFP; Muscle: QUAS:GFP; *mylz2*:QF2; Extracellular space: *ubb*:SEC-Rno.Ncan-EGFP), allowing for 3D segmentation and volume estimation for each of those fin compartments as well as overall fin volume across time and conditions. The first step was to extract the overall fin volume. For the Epithelia and Endoskeletal disk datasets, fin volume was quantified using the pipeline described above, using the signal of membranes labelled with claudinb:Lyn-GFP reporter. For the Extracellular space dataset, fin volume was quantified using the pipeline described above, using the signal of membranes labelled with the b-act:lyn-tdTomato reporter. For the Muscle dataset, fin volume was quantified using the pipeline described above, using the signal of membranes labelled with cellmask. The 3D fin volume masks were then applied to the channel with the fin compartment-specific label to obtain the compartment-specific signal restricted to the fin volume. For the Epithelia, another surface was generated on IMARIS using the parameters as above. The Edit/Selection/Delete tab was used to delete regions inside the pectoral fin since the *claudinb*:Lyn-GFP transgenic also labelled the membranes of some cells inside the fin. The resulting surface was used to generate an epithelial mask in which all voxels enclosed by the surface were retained and all voxels outside the surface were assigned a value of 0. As this mask generated by imaris contained holes in it and did not entirely encompass the epithelia, few more post processing steps were required. First, a gaussian smoothing operation of radius 10 μm was applied to ensure that a smooth mask was generated. Then, the mask was thresholded to create a binary mask of voxels with intensity greater than 1. Then a morphological closing operation of radius 3 μm was applied to generate the final binary mask of the Epithelia tissue. For the Endoskeletal disc, the masked tissue-specific channel was segmented using PlantSeg[23], a three-dimensional cell-segmentation pipeline, to generate an instance-segmentation image of the cartilage cells. The individual cell labels were combined into a single binary mask. A two-dimensional morphological closing operation with a radius of 2 μm was then applied independently to each XY slice to generate the final binary mask of the endoskeletal disc. For the Muscle and Extracellular space datasets, the masked tissue-specific channel was denoised using the median filter from the *clesperanto* package[24] with an x, y and z radius of 1. The image intensities were then normalized by dividing pixel values by the 99$^{th}$ percentile of the intensity of the entire image, and then rescaled to the 8-bit (0-255) range. To perform 3D segmentation for each of the fin's specific compartments, the Napari accelerated pixel and object classifier was used[24]. A different pixel classifier was trained for each of the fin tissue types. The classifier was trained manually on Napari by annotating the pixels belonging to the tissue and the pixels that were the background. The features that were used for training were gaussian blur, difference of gaussian, laplace box of gaussian blur and sobel of gaussian. The tissue-specific classifier was then applied to the tissue-specific channel to generate a prediction. The background label was deleted and the tissue label was retained to generate the binary masks for the Muscle and Extracellular Space. The final masks were then manually verified and corrected by hand if necessary. Specific fin compartment volume in $\mu m^3$ was quantified by multiplying the number of voxels inside the binary mask for each tissue by the acquired voxel dimensions. To obtain the normalized tissue volume across datasets, each specific fin tissue volume was divided by the respective overall fin volume obtained earlier. Statistical analysis and comparison between volumes of fin compartments was performed on python using the *statannot* package[22] by carrying out non-parametric, unpaired, two-tailed Mann Whitney *t*-tests between the different conditions. Statistical experimental details can be found in figure legends.

**Absolute and relative growth rate quantification**

To compute growth rates (Fig. 1G-H), we performed a non-parametric regression of fin volume as a function of time. For each condition, measurements were grouped by developmental time point and summarized by mean and standard deviation. We then fitted a Gaussian process (GP) regression model to the mean volume trajectory as a function of time, using a squared-exponential (radial basis function) kernel using the sklearn.gaussian_process function from the Python package, sklearn[25], with fixed time-scale of temporal smoothing equal to 50 h (which is close to the value of the function's auto-recommender). Experimental variability at each time point was incorporated as heteroscedastic observational noise, using the measured standard deviation as the noise level. From the posterior distribution of the fitted Gaussian process, we drew multiple samples of smooth volume trajectories over a dense time grid. The instantaneous volume growth rate $dV(t) / dt$ was computed numerically as the time derivative of each sampled trajectory. The relative growth rate was obtained as $[dV(t)/dt] / V(t)$. Reported growth curves correspond to the median across posterior samples, and uncertainty bands indicate the 16%–84% percentiles (i.e., approximately ±SD). This non-parametric approach avoids assuming a specific functional form for growth dynamics while providing smooth derivatives and principled uncertainty estimates.

**Fin 2D midsurface generation, area and thickness quantification**

Curvilinear 2D fin midsurfaces were obtained using a custom-made computational pipeline (Python) that automatically generates a two-dimensional midsurface in the center of the relatively flat fin and assigns a curvilinear coordinate system. As input, this pipeline takes the three-dimensional fin volume binary masks and a manually annotated direction of the proximal-distal axis (PD), together with a preliminary estimate of the direction of the anterior-posterior axis (AP) per fin. In the first step, a planar cross-section of the 3D volume mask through its center-of-mass and perpendicular to the preliminary anterior-posterior axis is used (Fig. S2A). In this cross-section, the algorithm determines a curved midline that has maximal distance from the boundaries of the cross-section. Specifically, the algorithm computes the midpoints of line segments defined as the intersection of the planar cross-section and coplanar rays along preliminary estimates for a local dorso-ventral axis that emanate from a manually provided reference point close to the center of the osculating circle of the curved proximal region of the midsurface cross-section (Fig. S2B). The user can then apply manual corrections, which might be needed, e.g., near the most proximal part of the fin. Next, multiple orthogonal cross-sections are computed as follows: at equidistant points along the midline, we compute the tangent vector along the midline and determine the plane normal to it. The intersection of this normal plane and the binary fin mask defines a family of orthogonal cross-sections. For each of these orthogonal cross-sections, we compute the midpoints of line segments parallel to the local normal vector of the midline (Fig. S2C). The set of these midpoints from all the orthogonal cross-sections defines a point cloud that lies in the middle between the dorsal and ventral surface of the three-dimensional fin volume, and allows to compute a smooth midsurface. For this, we use the Python library *pyvista*[26], obtaining a triangulated mesh with a spatial resolution of about 1-5 µm (Fig. S2D). Finally, the nodes of this triangulated mesh are scaled from voxel coordinates to micrometers using the known voxel size. The *fin area, A,* is then computed as the total area of the triangulated midsurface mesh. Python code is available at: https://github.com/Coolix99/ZF_PF_Geometry. To make sure that surfaces generated from different transgenic backgrounds could be compared across datasets, we imaged fish carrying both ubiquitous nuclei and membrane labels and understood that the different labels used render different volumes and consequently thicknesses, but fin area is comparable across the two signals (Fig. S3G).

**Alternative computation of fin thickness**

To assess the robustness of our quantification of fin thickness, we additionally computed a local fin thickness $h(x_{PD},x_{AP})$ at every point of the fin midsurface, analogous to the computation of $L_{DV}$ at the reference point $(x_{PD},x_{AP})=(40\%\ L_{PD},\ 0)$. We computed the spatial mean of the local thickness (for this, we first computed the local thickness at the midpoint of each triangle of the triangulated midsurface and then computed the weighted average of these local thicknesses, weighted by triangle area). We confirmed that this mean thickness agrees with the ratio $V/A$ up to numerical precision (compare Fig. S3A with Fig. S3C). This alternative mean thickness also follows a very similar trend to $L_{DV}$ as a function of time (Fig. 3B), in both injured and control fins.

**Curvilinear coordinate system of fin midsurface and axes length**

We define a *curvilinear coordinate system* on the fin midsurface with coordinates ($x_{PD}$,$x_{AP}$) as follows.At the beginning, the user manually annotated the proximal-distal direction, represented by a unit vector $\mathbf{e}_{PD}^{(0)}$. This constant vector $\mathbf{e}_{PD}^{(0)}$ defines local unit vectors $\mathbf{e}_{PD}$ for each triangle of the midsurface by simple projection. Thus, at each point $\mathbf{r}_i$ of the triangular mesh, we have three unit vectors: the position-dependent vector $\mathbf{e}_{PD}$ pointing along the local proximal-distal axis, the position-dependent normal vector of the triangle $\mathbf{n}_i$, and a vector $\mathbf{e}_{AP} = \mathbf{n}_i \times \mathbf{e}_{PD}$ pointing along the local anterior-posterior axis (Fig. S2D). Integrating the unit vectors $\mathbf{e}_{PD}$ and $\mathbf{e}_{AP}$ along the triangulated surface defines corresponding coordinates $x_{PD}$ and $x_{AP}$ in proximal-distal and anterior-posterior direction, respectively. Note that these coordinates are at first only determined up to an arbitrary offset. This curvilinear coordinate system defines a (non-isometric) mapping from the midsurface embedded in three-dimensional space onto a two-dimensional representation spanned by the curvilinear coordinates $x_{PD}$ and $x_{AP}$ (Fig. S2E). This cartographic 2D-projection will be integral to average spatial quantities across individuals (see section: 'Proliferation density maps'). To fix the yet undefined offset of the curvilinear coordinates $x_{PD}$ and $x_{AP}$, we consider the bounding box of the cartographic 2D-projection of the fin midsurface, with the edges of the bounding box parallel to the coordinate axes. Specifically, this bounding box spans across [min($x_{PD}$), max($x_{PD}$)] along the proximal-distal direction and across [min($x_{AP}$), max($x_{AP}$)] along the proximal-distal direction, where the maximum/minimum is taken over all points of the 2D-projected midsurface. Without loss of generality, we can require min($x_{PD}$) = 0, max($x_{PD}$) = $L_{PD}^{BB}$, -min($x_{PD}$) = max($x_{PD}$) = $L_{AP}^{BB}$ / 2, where we introduced the dimensions of the bounding box $L_{PD}^{BB}$ = max($x_{PD}$) - min($x_{PD}$) and $L_{AP}^{BB}$ = max($x_{AP}$) - min($x_{AP}$), which fixes the offset of the coordinates.

The 2D-projection of the midsurface allows defining characteristic lengths for each fin along the developmental proximal-distal (PD), anterior-posterior (AP) and dorsal-ventral (DV) axes. For this, we use a reference point in the 2D-projection located at $x_{PD}$ = 40% $L_{PD}^{BB}$, xAP = 0 (Fig. S2E), together with the pre-image of this reference point in the three-dimensional fin mask under the cartographic projection. We consider the lengths of curvilinear coordinate lines passing through this reference point. Note that the cartographic projection from the fin midsurface embedded in three-space onto its 2D-projection is not isometric, i.e., generally length measurements differ depending on whether these are conducted on the curved midsurface or the 2D-projection. However, our definition of the curvilinear coordinate system ensures the cartographic projection preserves lengths along coordinate axes, hence the length measurements that define $L_{PD}$ and $L_{AP}$ below are independent of whether these are conducted on the curved midsurface or the 2D-projection. For the characteristic length $L_{PD}$ along the proximal-distal axis, we compute the length of a line along the proximal-to-distal direction passing through the reference point, i.e., the midline (Fig. S2E). For the characteristic length $L_{AP}$ along the anterior-posterior direction, we similarly compute the length $L_{AP}$ of a line along the anterior-posterior direction, again passing through the reference point. For the characteristic length $L_{DV}$ along the dorso-ventral axis, finally, we compute in the three-dimensional fin mask the length of a line along the dorso-ventral axis normal to the fin midsurface passing through the pre-image of the reference point.

Statistical analysis and comparison between fin axes lengths was performed using either GraphPad Prism or Python using the statannot package[22] by carrying out non-parametric, unpaired, two-tailed Mann Whitney *t*-tests between the different conditions. Statistical experimental details can be found in figure legends.

**Fin anisotropy and aspect ratio**

Fin anisotropy ε was quantified from the PD and AP length values ($L_{PD}$, $L_{AP}$) from the live imaging datasets, as described previously[4]. Anisotropy is defined as the ratio of the growth rates $g$ along the AP versus PD axis $\varepsilon = (g_{AP}/g_{PD})$, where $g_{AP} = (dL_{AP}/dt \;/\; L_{AP})$ with $dL_{AP}/dt$ the time derivative of the $L_{AP}$ length and $g_{PD} = (dL_{PD}/dt \;/\; L_{PD})$. Therefore, $(dL_{AP} \;/\; L_{AP}) = \varepsilon\, (dL_{PD} \;/\; L_{PD})$ and taking integrals $\log L_{AP} = \varepsilon \log L_{PD} + C$, where C is an integration constant. Thus, we estimated growth anisotropy ε by fitting to the length data the power-law relationship $L_{AP} \sim L_{PD}^{\varepsilon}$ and displaying the relationship as a log-log plot where the slope of the line corresponds to the anisotropy.

Aspect ratio was quantified by plotting the obtained PD versus AP length values ($L_{PD}$, $L_{AP}$) or PD versus DV length values ($L_{PD}$, $L_{DV}$). Statistical analysis and comparison between fin axes aspect ratios was performed on python using the statannot package[22] by carrying out non-parametric, unpaired, two-tailed Mann Whitney *t*-tests between the different conditions. Statistical experimental details can be found in figure legends.

**Cell segmentation, number and proliferation quantification**

Nuclei segmentation was performed using a custom machine learning code, closely following methods in Cellpose[27] and Omnipose[28]. First, we manually annotated cropped ROI from the nuclei channel from 3D image z-stacks from H2B-mCherry; mAG-zGeminin double transgenics. Using this ground truth data, a convolutional neural network was trained on predicting a flow field towards the center of each nucleus, as well as a binary mask of the nuclei. In the following segmentation step, the position of each pixel in this binary mask is propagated along the flow field. This displaced point cloud is clustered using DBSCAN[29]. By measuring our data, we considered that all segmented objects larger than 22 $\mu m^3$ represent nuclei. This defines the nuclei number. Next, all nuclei are classified as either zGeminin-positive or negative. For this a random forest classifier was trained on manually annotated ground truth data, where equal rates of false positive and false negative classifications for a test data set were imposed. For the classifier, three features were used, namely average mAG-zGeminin intensity in the nucleus region, a nucleus core region obtained by morphological erosion of 2px, and a region including neighboring pixels, obtained by morphological dilation of 5px. By dividing the total segmented nuclei per fin by the segmented mAG-zGeminin+ nuclei, we obtain the reported fraction of mAG-zGeminin positive cells across conditions (Fig. 2J).

**Relationship between proliferation rate and cell number**

By assuming a constant time duration of the zGeminin+ phase, $t_{Geminin}$, the relative fraction zGeminin+ cells can be converted into a proliferation rate by dividing by this time. Integration of this proliferation rate in time yields a model prediction for the cell number $N(t)$ as a function of time, where the time duration $t_{Geminin}$ of the zGeminin$^+$ phase (S/G2/M[6]) was used as fit parameter (Fig. 2K, dashed curves). Specifically, we use the total number of segmented nuclei in the H2B-mCherry; mAG-zGeminin double transgenics data as a proxy for the total cell number $N(t)$, which is reported in Fig. 2K. We further assume that the fraction of zGeminin+ cells in this data represents a proxy for the proliferation rate, which is reported in Fig. 2J. Specifically, we determined the mean fraction $p_{Geminin+}(t)$ of zGeminin+ cells by averaging the local fraction of mAG-zGeminin positive nuclei across the entire fin. For the control condition of development, we fit an exponential decay to this time-dependent proliferation rate (Fig. 2J, dashed curve, decay time = -0.03371, $R^2$=0.89). To convert the fraction of proliferating cells to a relative proliferation rate $g_N(t)$, we divided this fraction by a constant duration of the zGeminin$^+$ phase (S/G2/M phase of the cell cycle, $t_{Geminin}$≈3.25h, Fig. S1E-G) according to $g_N(t) = p_{Geminin+}(t) / t_{Geminin}$. We can then computationally reconstruct the expected cell number $N(t)$ at time t by integrating the following equation $d\,N(t) / dt = g_N(t)\,N(t)$ . The results of this simple growth model are presented as dashed curves in Fig. 2K; the initial cell number $N(t = 48\text{ h})$ was determined by a fit using the time duration as $t_{Geminin}$ as an additional fit parameter. The fitted time duration of $t_{Geminin}$≈3.25 h is comparable yet shorter than our direct measurements (~5h, Fig. S3E-G).

**Cell cycle measurements**

Using the acquired light sheet datasets of mAG-zGeminin; H2B-mCherry double transgenic larvae, obtained z-stacks were converted into HDF5 format using the BigDataViewer plugin in Fiji[30]. From these, individual nuclei were manually tracked using the Mastodon Fiji plugin[31]. S/G2/M phases of the cell cycle were tracked using the mAG-zGeminin channel, while labelling all nuclei (in H2B-mCherry channel). Cell cycle phases were defined as: (i) S/G2/M, the interval of time encompassing the start of mAG-zGeminin expression until the positive cell undergoes cytokinesis; (ii) G1, the interval of time since a cytokinesis event until daughter cells start expressing mAG-zGeminin; and (iii) full cell cycle, the combined intervals of (i) and (ii), i.e. the interval of time in which a single cell starts expressing mAG-zGeminin, it divides, its daughter cells undergo G1 and start re-expressing mAG-zGeminin, until their division. The length of each phase was determined from Mastodon tracks. Results were plotted using GraphPad Prism, displaying the individual data points with mean ± SD.

**Proliferation density maps**

Fig. 2I reports the position-dependent fraction of zGeminin$^+$ cells, averaged over several individuals. For this, we developed a mapping of individual fin midsurfaces to a reference shape of the fin, according to its developmental stage. In short, all center points of nuclei were first projected onto the individual fin midsurface (Fig. S2D), together with their classification as zGeminin$^+$ or zGeminin$^-$ nuclei. These projected nuclei centers were then mapped from their individual fin midsurface to an average reference shape, ω. This allowed us to pool the classified nuclei centers from all individuals and compute a sample-averaged local fraction of zGeminin+ cells on the respective reference midsurface.

To compute ω for every condition and every developmental time point, we “averaged” the boundaries of the cartographic projections of individual fin midsurfaces as detailed below. Then cartographic projections of individual fin midsurfaces were mapped onto ω by first matching their boundaries and then extending this mapping to the interior such as to minimize distortions, similar to minimizing an elastic deformation energy. Specifically, each nucleus is assigned to the nearest node of the three-dimensional midsurface. As the curvilinear coordinate system ($x_{AP}$, $x_{PD}$) defines a two-dimensional cartographic projection, we can represent the midsurface in two dimensions (Fig. S2E). To average between individuals, we compute for each group (experimental condition and developmental time point) an average fin boundary b($s$) parametrized by arclength $s$ by averaging the first elliptic Fourier modes of the boundary $B_j(S)$ of the cartographic projections of the fin midsurfaces of the respective individuals (labeled $j$). We define a one-to-one mapping between each $B_j(S)$ to b($s$) by identifying the most distal point of each $B_j(S)$ as well as b($s$), and linearly interpolate along the boundary. To find a mapping not just of the boundaries but also of the interiors, we calculate the displacement field u=x-X between the coordinates of an individual boundary $X \in Bj$ and those of the average boundary $x \in b$. By solving $\Delta u=0$ for all points $X \in \Omega$ in the interior of $Bj=\partial\Omega j$ and keeping $u(X|Bj)$ fixed, we find a smooth interpolation of the boundary displacement, mapping $\Omega j$ to $\omega$ with $\partial\omega=b$. This mapping can be interpreted as minimizing a pseudo-elastic deformation energy, corresponding to a linear elastic material with fixed boundary displacement. This mapping of a reference shape enables mapping every node of the triangulated mesh representation of individual fin midsurfaces, together with any associated data, to the average shape ω. On this average shape, one can now introduce a triangulation of possibly lower spatial resolution, whose nodes serve as bin centers of a two-dimensional histogram. Thus, every nucleus in an individual fin is mapped to the nearest node of the average fin midsurface given by ω. Pooling this data, we calculate the local fraction of zGeminin$^{+}$ cells.

**Immunostainings signal quantification**

*Per fin volume:* 3D pectoral fin volume masks based on the nuclei signal were generated in Imaris (version 10.1.1) as described above, and used to extract the total Smoc1 or Phospho-Smad1/5/9 signal intensity within the fin for each sample. To remove high background levels resulting from Phospho-Smad1/5/9 immunostainings, a processing pipeline adapted from [4] was generated as a Python custom script. Briefly, bright hot pixels in the pSmad1/5/9 channel were removed using a Fiji-style *Remove Outliers*[21] filter (radius = 3 px, threshold = 200), reimplemented in Python (scipy.ndimage): pixels exceeding the local median by more than the threshold specified were replaced with that median, computed per z-slice. Filtering was applied only to the pSmad1/5/9 channel; all other channels were unmodified. Then 3D pectoral fin volume masks based on nuclei signal (DAPI) were generated in Imaris (version 10.1.1) as described above, and used to extract the mean Phospho-Smad1/5/9 signal intensity within each fin mask. To compare pSmad1/5/9 values across fins of different volumes, we normalized individual intensity by an arbitrary volume, 100 $\mu m^3$ (Fig. 4K), followed by plotting the individual data per dataset and timepoint, jointly with respective mean and SD.

*Per region of interest (ROI):* After image acquisition, 3D rotation, cropping and flattening was performed using IMARIS, so that the fin's endoskeletal disc was aligned along the AP/PD plane. Maximum intensity projections were obtained and ROIs were defined using Fiji[21]. To characterize *Smoc2*$^{-/-}$ mutants (Fig. S6B-D), a ROI midline was defined by an area of 50 pixels width in the endoskeletal disc, along the region abutting the fin fold, which was identified by nuclear density monitored with DAPI signal, as performed in [4]. The signal intensity of Smoc1 or

Phospho-Smad 1/5/9 in a particular position along the ROI midline corresponds to the average signal in their orthogonal positions within the ROI. To characterize the distribution of Smoc1 along the proximal-distal axis of the fin (Fig. S5A-B), a ROI line was defined by an area of 50 pixels width along the proximal-distal axis of the fin. As before, the signal intensity of Smoc1 corresponds to the average signal in their orthogonal positions within this ROI. In both cases, individual profiles were normalized by dividing ROI length values by respective maxima, followed by interpolation, averaging and plotting mean together with SEM.

**Sequence alignment**

Sequences for DrSmoc1 (GenBank, MK285359)[4]; DrSmoc2 (Uniprot, A0AC58H334 - isoform X1); DrBMP2a (Uniprot, O13109); DrBMP2b (Uniprot, O93369); DrBMP4 (Uniprot, O57574); HsSMOC1 (UniProt, Q9H4F8 - canonical); HsSMOC2 (UniProt, Q9H3U7 - canonical); HsBMP2 (UniProt, P12643), HsBMP4 (UniProt, P12644), CeDBL-1 (UniProt, G5EEL5) were obtained and pairwise alignments were made using Biopython[32] Bio.Align.PairwiseAligner package. Specifically, we employed a BLOSUM62 substitution matrix[33] and used a global, full-length Needleman-Wunsch-style alignment mode; had gap penalties with parameters affine, open 10, and extend 0.5; and applied an anchor/star alignment strategy, where each block picks one fixed anchor sequence, and every other sequence in that block is independently pairwise-aligned against only that anchor; sequences were never aligned against each other directly, and there was no joint multi-way optimization. We used three anchors:
TY1 block - DrSmoc1 residues 138–183
TY2 block - DrSmoc1 residues 299–346
BMP block - DrBmp2a mature-domain sequence

***In silico* protein-protein interaction structure predictions**

The ColabFold v1.6.1[34] implementation of AlphaFold2-multimer_v3[35,36] was used to predict structures of complexes involving combinations of one copy of the secreted form of DrSmoc1 and two copies of the mature form of DrBMP2a (known to homodimerize). Parameters used were: MMseqs2 MSA mode, no templates, 3 recycles, 5 models, rank-1 Amber-relaxed. A 5Å heavy-atom contact-distance cutoff was used to build the 150-contact table. The result was a structural prediction in which DrSmoc1 interacts with DrBMP2a in a 1:2 stoichiometry, with an interface predicted template modeling (ipTM) score of 0.80 (pTM 0.57, mean pLDDT 73.4, max PAE 31.7Å). This is in agreement with published structural complexes in *H. sapiens*, where SMOC1:BMP2:BMP2 complexes have a ipTM score of 0.81[3]. The ipTM score is a confidence score generated by AlphaFold[35]. 3D rendering was performed using 3Dmol.js[37].

### Supplementary Theory Notes

##### 1. A simple timer model

We start by presenting a simple timer model, which will turn out to be insufficient to explain the experimental growth data. In this model, the relative growth $g(t)$ follows a fixed, programmed time course, and does not dynamically adapt to current fin size or injury. Thus, this model corresponds to open-loop control without feedback.
The fin area $A(t)$ evolves according to

(1) $\mathrm{d}A(t)/\mathrm{d}t = g(t)\, A(t)$ .

We choose the time-dependent growth rate $g(t)$ such that the model exactly reproduces the dashed theory curve for the control condition shown in Fig. 3I, starting from an initial fin area $A(t_0) = A_0 = 1.8\cdot10^4$ µm$^2$ with start time $t_0$=48 hpf.

Eq. (1) can be solved by direct integration:

$\mathrm{A}(t) = \gamma(t)A_0$ with $\gamma(t) = \exp[G(\mathrm{t})]$,
where
$G(t) = \int_{t0}^{t} g(t)\, \mathrm{d}t$

is the cumulated relative growth. Thus, $\gamma(t)$ denotes the factor by which the fin area has increased between $t_0$ and time $t$.

By construction, the timer model exactly reproduces the mean dynamics for control conditions. However, the model fails to reproduce the growth adaptation observed for the different injury conditions (Fig. 3G, upper panel). Because all fins follow the same prescribed growth protocol, their areas are multiplied by the same factor $\gamma(t)$, irrespective of the amount of tissue removed. In particular, all fins grow by the same factor $\gamma(t$=144 hpf) between 48 hpf and 144 hpf.

The timer model also cannot reproduce the experimentally observed reduction in relative size variability, as quantified by the coefficient-of-variation of fin area (Fig. 3G, lower panel). To see this, assume that the initial fin size is not equal to a constant value $A_0$ but distributed according to a distribution $p_0(A)$ with mean $<\mathrm{A}(t_0)> = A_0$ and standard deviation $\sigma_0$. The initial coefficient of variation is thus CV = $\sigma_0/A_0$. The distribution of areas at time $t$ is given by $p(A,t) = p_0(A\, /\, \gamma(t))\, /\, \gamma(t)$, which has mean $<A(t)> = \gamma(t)A_0$ and standard deviation $\sigma(t)=\gamma(t)\sigma_0$. Thus, the coefficient of variation stays constant in time $\mathrm{CV}(t) = \sigma(t)\, /\, A(t) = \sigma_0\, /\, A_0 = \mathrm{CV}(t$ =48 hpf), see Fig. S4A. Analogous arguments are well established in theories of cell-size control, where size-independent timer mechanisms fail to provide size homeostasis under exponential growth[38]. Thus, a fixed timer program can neither account for injury-dependent growth adaptation nor reduce relative differences in fin size.

#### 2. *Integral-feedback control of area growth*

As a minimal model for the growth dynamics of the pectoral fin, we propose a set-point model for fin area, $A$. As in Eq. (1), we assume that the change of the fin area d$A$/d$t$ is given by a relative area growth rate $g(t)$, such that $g(t)$. Yet, now we assume that $g(t)$ is dynamically controlled by fin area $A$. The simplest assumption of instantaneous feedback, $g(t)=g(A(t))$, is incompatible with the observation that initially, after the amputation, proliferation and therefore growth is delayed. Therefore, we model the growth rate with a delay

(2) $\tau\, dg/dt = g_{target}(A) - g$ ,

where \tau denotes a feedback time scale on which tissue growth reacts to changes in tissue size. The simplest equation for $g_{target}(A)$, allowing for growth arrest at a target size, is

(3) $g_{target}(A) = g_0 ( 1 - A/A_{target} )$ ,

where $g_0$ sets a characteristic growth rate and $A_{target}$ denotes the size setpoint of the fin. For $A > A_{target}$, the target growth rate can become negative, i.e., the fin can (slightly) shrink.

This minimal model can be regarded as a variant of integral-feedback control[39,40]. Indeed, the current growth rate $g(t)$ accumulates the past size-dependent error signal $g_{target}(A)$, similar to an integral-feedback controller, though as a leaky integrator with memory timescale \tau). The model exhibits perfect adaptation because at steady state, we must have $g$=0 (by Eq. (1)), hence $g_{target} = 0$ (by Eq. (2)), hence $A = A_{target}$ (by Eq. (3)). Moreover, its feedback reduces relative size variability, see Fig. S4B.

The proposed model has three model parameters \tau, $g_0$, and $A_{final}$; as well as two initial conditions $A(t_0)$ and $g(t_0)$. The initial growth rate is set to its steady-state value for development scenarios, yet is set to zero for regeneration scenarios, reflecting the growth pause following the wound response. We can successfully fit all experimental scenarios using a common set of model parameters (\tau, $g_0$, and $A_{target}$), assuming only this different initialization of initial growth rate in injured fins and different initial fin sizes $A(t_0)$, see Fig. 3I and Fig. 4H. The only exception is the uninjured control condition for the Smoc1$^{-/-}$;Smoc2$^{-/-}$ mutant, where a reduced value of $A_{target}$ was assumed based on the experimental data (Fig. 4F).

We can successfully fit the growth curves for uninjured control and all three injury conditions shown in Fig. 3I with a common set of model parameters (\tau, $g_0$, and $A_{final}$), assuming only different initial conditions. For Fig. 4H, we then re-use the model parameters from Fig. 3I, with the exception of the reference area $A_{target}$ for the uninjured control condition in the Smoc-/- mutant. We perform these shared fits with a Bayesian approach, allowing us to obtain a probability distribution of the model parameters from our experimental observations. As model priors, we used log-normal distributions as specified in the table below. The likelihood model (mimicking experimental variability and measurement noise) was chosen as normal distribution
$A_{experiment} \sim \mathrm{Normal}( A_{model}, \sigma = \varepsilon_{abs} + A_{model}\, \varepsilon_{rel})$,

where the absolute and relative errors $\varepsilon_{abs}$ and $\varepsilon_{rel}$ are hyperparameters, which are automatically fitted. We used a multi-chain Monte-Carlo method to sample parameters from the posterior distribution using the software tool *stan* (https://mc-stan.org/).

Mean model parameters as reported in Table S2 correspond to mean values of marginalized posterior distributions. The solid curves shown in Fig. 3I and Fig. 4H correspond to those parameters.

Model priors:

| **Parameter** | **Prior distribution** |
|---|---|
| $g_0$ | $\log_{10}(g_0/h^{-1}) \sim \text{Normal}(-1.1, 0.5^2)$ |
| $\tau$ | $\log_{10}(\tau/h) \sim \text{Normal}(0.5, 0.5^2)$ |
| $A_{target}$ | $\log_{10}(A_{target}/10^4\ \mu m^2) \sim \text{Normal}(1.0, 0.15^2)$ |
| $A(t_0 = 48\ \text{hpf})$ | $\log_{10}(A(t_0 = 48\ \text{hpf})/10^4\ \mu m^2) \sim \text{Normal}(0.3, 0.15^2)$ |
| $A(t_0 = 72\ \text{hpf})$ | $\log_{10}(A(t_0 = 72\ \text{hpf})) \sim \text{Normal}(0.75, 0.15^2)$ |
| $\varepsilon_{abs}$ | $\varepsilon_{abs}/10^4\ \mu m^2 \sim \text{HalfNormal}(0, 2^2)$ |
| $\varepsilon_{rel}$ | $\varepsilon_{rel} \sim \text{HalfNormal}(0, 0.1^2)$ |

We chose area $A$ instead of volume $V$ as control variable for three reasons: (i) Fin area converged to approximately the same final fin area at 144 hpf for all injury conditions and the uninjured control, suggestive of an area setpoint (Fig. 3C, upper panel). In contrast, final fin volume was more variable across the different injury conditions (Fig. S3F). (ii) We observe a strong reduction in the coefficient-of-variation of fin area (Fig. 3G, lower panel), suggestive of a feedback mechanism that reduces relative area variations. In contrast, for fin volume, the reduction in the coefficient-of-variation was less pronounced (Fig. S3F). (iii) Growth along the dorso-ventral axis contributes to volume growth, suggesting a partial decoupling from area (Fig. 3C). Moreover, area growth appears to be characterized by a coupling between the proximal-distal and anterior-posterior axes, with a stereotypic in-plane anisotropy $\varepsilon$ of growing fins that is rapidly restored after injury (Fig. 3D, Fig. S3B). This stereotypic in-plane anisotropy allows to approximately express the length-scales $L_{PD}$ and $L_{AP}$ as a function of the fin area $A$ as $L_{PD} \sim A^{1/1+\varepsilon}$ and $L_{AP} \sim A^{\varepsilon/1+\varepsilon}$, further motivating a model that focuses on area growth, as opposed to growth along the individual PD and AP axes.

#### Supplementary Tables

Table S1. Sequences of primers used qPCR analysis during this study.

| Gene | Accession # | Forward primer (5’) | Reverse primer (3’) |
|---|---|---|---|
| *ef1α, eef1a1l1* | NM_131263 | AAATGGCCAAACAAGGG AACACG | TCCAATAGAGTAACACCA CCGGC |
| *smoc2* | XM_073920286.1 | GTCGAGGGCAATGCAAA GATAC | TCAGGATTGCAGACAGG CAC |
| *smoc1* | NM_001201393.2 | GTTGACCGGTATGAAAGA AGCAG | CAAATGTGCCGTCGTCGT TG |

Table S2. Fit parameters for integral-feedback control growth model.

| Parameter | mean | SD | |
|---|---|---|---|
| $A_{target}$ (same for all conditions, except one) | 11.2 | 0.2 | $10^4$ μm$^2$ |
| $A_{target}$ (Smoc$^{1-/-}$ Smoc$^{2-/-}$ , uninjured control) | 8.3 | 0.2 | $10^4$ μm$^2$ |
| τ (same for all conditions) | 7.3 | 1.0 | h |
| $A_0$ (wildtype, uninjured control) | 1.8 | 0.1 | $10^4$ μm$^2$ |
| $A_0$ (wildtype, 30% cut) | 1.3 | 0.1 | $10^4$ μm$^2$ |
| $A_0$ (wildtype, 50% cut) | 1.0 | 0.1 | $10^4$ μm$^2$ |
| $A_0$ (wildtype, 30% cut at 72 hpf) | 4.7 | 0.2 | $10^4$ μm$^2$ |
| $A_0$ (Smoc$^{1-/-}$ Smoc$^{2-/-}$, uninjured control) | 1.7 | 0.1 | $10^4$ μm$^2$ |
| $A_0$ (Smoc$^{1-/-}$ Smoc$^{2-/-}$, 30% cut) | 0.8 | 0.0 | $10^4$ μm$^2$ |
